# Large-Scale, Multi-Year Microbial Community Survey of a Freshwater Trout Aquaculture Facility

**DOI:** 10.1101/2022.05.03.490559

**Authors:** Todd Testerman, Lidia Beka, Stephen R. Reichley, Stacy King, Timothy J. Welch, Gregory D. Wiens, Joerg Graf

**Author notes:** Lidia Beka, National Cancer Institute, National Institutes of Health, Bethesda, Maryland, USA. Stephen R. Reichley, College of Veterinary Medicine and Global Center for Aquatic Health and Food Security, Mississippi State University, Starkville, Mississippi, USA. Stacy King, College of Veterinary Medicine & Biomedical Sciences, Veterinary Diagnostic Laboratories, Colorado State University, Fort Collins, Colorado, USA.

## Abstract

Aquaculture is an important tool for solving growing worldwide food demand, but infectious diseases of the farmed animals represent a serious roadblock to continued industry growth. Therefore, it is essential to understand the microbial communities that reside within the built environments of aquaculture facilities to identify reservoirs of bacterial pathogens and potential correlations between commensal species and specific disease agents. Here, we present the results from three years of sampling a commercial rainbow trout aquaculture facility. The sampling was focused on the early-life stage hatchery building and included sampling of the facility source water and outdoor production raceways. We observed that the microbial communities residing on the abiotic surfaces within the hatchery were distinct from those residing on the surfaces of the facility water source as well as the production raceways, despite similar communities in the water column at each location. Within the hatchery building, most of the microbial classes and families within surface biofilms were also present within the water column, suggesting that these biofilms are seeded by a unique subgroup of microbial taxa from the water. Lastly, we detected a common fish pathogen, *Flavobacterium columnare*, within the hatchery, including at the source water inlet. Importantly, the relative abundance of this pathogen was correlated with clinical disease. Our results characterized the microbial communities in an aquaculture facility, established that the hatchery environment contains a unique community composition, and demonstrated that a specific fish pathogen resides within abiotic surface biofilms and is seeded from the natural source water.

**Importance:** The complex microbial consortium residing in the built environment of aquaculture facilities is poorly understood. In this study, we provide a multi-year profile of the surface- and water-associated microbial communities of this biome. The results demonstrated that distinct community structures exist in the water and on surfaces. Furthermore, it was shown that a common and economically impactful bacterial pathogen, *F. columnare*, is continually introduced via the source water, is widespread within surface biofilms in the hatchery environment, and is likely amplified within these raceways but does not always cause disease despite being present. These results advance our understanding of pathogen localization at fish farms, show the interplay between host and environmental microbiomes, and reveal the importance of microbial community sequencing in aquaculture for identifying potential beneficial and harmful microbes. This study adds to the aquaculture microecology dataset and enhances our ability to understand this environment from a “One Health” perspective.

## Introduction

Aquaculture has grown extensively over the past three decades, but disease caused by microbes poses one of the greatest threats to the continued growth of this industry (1). While the microbiomes of many farmed fish have been studied (2, 3), the microbial communities existing within the built environments of these animals, including raceway surfaces, have garnered less attention. These biomes are important as they offer potential insight into the distributions and abundances of specific pathogens. For example, in farms where untreated surface water is used, microbes from local upstream ecosystems flow into raceways and form biofilms on their surfaces (4). A mature biofilm is typified by a sturdy and stable architecture, a protective matrix, and often, a highly dense and heterogenous assembly of microbial species. These features allow biofilms to serve as microbial banks, providing stable sites for genetic exchange that are relatively undisturbed by environmental hazards, including antibiotic and chemical treatments (5). Previous studies have demonstrated that most biofilm-associated microbes in aquaculture facilities are not harmful, and many may in fact be beneficial to the farmed fish. However, a subset of species are opportunistically pathogenic and can cause rapid and deadly disease outbreaks depending on host health, age, and environmental factors (6, 7).

Numerous methods are used to farm rainbow trout (*Oncorhynchus mykiss*) in the United States, with a common approach being the flow-through system in which natural spring or river water is diverted through the facility and eventually flows out (8). This type of system can provide relatively stable temperatures and consistent water quality based on the farm’s location, source water type, flow, and distance the water travels above ground level (8). These farms are typically configured such that eggs and young fish are housed within a hatchery building, or early life-stage rearing facility, where they mature for several months (9). After reaching a specified size, fish are moved into outdoor raceways for grow-out before harvesting. Although disease can occur at any step in this process, generally the younger individuals are at greatest risk relative to the older individuals. Due to an immature immune system and difficulties with vaccination, rapid and significant mortality events occur often in the hatchery setting (10).

Two bacterial species, *Flavobacterium columnare* and *Flavobacterium psychrophilum*, have been shown to impact the early life stages of farmed rainbow trout. *F. columnare* causes columnaris disease (CD) (11) and has recently been divided into four species based on genomic differences, with each species being associated with varying virulence levels in laboratory studies (12). *F. psychrophilum* causes bacterial cold-water disease (BCWD), which is often observed in immature cold water fish species (13). There is only one licensed vaccine for CD in catfish (14), and no licensed vaccines are available for BCWD within the United States (although a live-attenuated vaccine is in development (15–17). The lack of potent, universal, and well-tolerated vaccines for fish, especially for the youngest fish (fry), means that early and accurate detection of pathogens is essential for reducing the impacts of an outbreak. Current mitigation and prevention methods heavily rely on animal health surveillance targeting fish that exhibit symptoms of CD or BCWD (i.e., lethargy, abnormal swimming, lack of appetite, and skin lesions) for culturing and diagnostic PCR. If a pathogen is identified, intervention efforts commence. However, while important, this strategy focuses on identifying pathogenic *Flavobacterium* spp. during the period of clinical disease, when bacterial load is high and peak shedding occurs, overlooking initial infection events, the incubation period, the window of subclinical disease, asymptomatic shedding, and importantly, the point source.

Outbreaks of *F. psychrophilum* and *F. columnare* frequently re-occur in the hatchery environment between different production lots despite disinfection procedures. Such observations suggest that disinfection protocols are not entirely effective at removing these pathogens or that they are regularly re-introduced into this environment. Both of these flavobacterial pathogens have been shown to survive for long periods of time in freshwater at cool temperatures in the absence of their fish host (18, 19). They have also been shown to readily form biofilms, which may increase their ability to persist and survive treatment with some of the common disinfectants used in aquaculture (20, 21). Understanding the reservoirs and sources of these pathogens will be an important step in developing management practices for their control.

High throughput sequencing of the 16S rRNA gene in samples collected at various sites within raceways and the inflowing water provides crucial insights into the sites of pathogen localization within farms along with potential correlations with collected metadata. Coupled with standard health surveillance and culturing strategies, microbiome research at fish farms allows us to characterize the transient versus resident flora of the farm, identify possible interactions between microbiota, and direct the development of new methods of pathogen control by identifying beneficial microbes for probiotic studies. Microbiome research at fish farms can thus inform and enhance outbreak control measures.

In this work, we present a study of the microbiome of a rainbow trout facility spanning three consecutive years of sampling. We performed sequencing of a hypervariable region in the 16S rRNA gene to characterize the native microbial communities flowing into and out of the hatchery raceways as well as those growing within biofilms formed on structures in the raceways. We sequenced water and surface samples collected from three sites: Box Canyon Spring (BCS), the hatchery facility, and the outdoor raceways, with a primary focus on the hatchery facility. We also aimed to leverage our 16S rRNA survey data to simultaneously evaluate the presence of two pathogenic flavobacterial species and better understand pathogen refuge and transmission within the hatchery.

## Materials and Methods

### Sample Sites and Site Metadata

Samples were collected from and around a rainbow trout aquaculture facility in Idaho, USA. Sites included BCS (where the source water is obtained), the hatchery building, and the outdoor raceways. Collection was performed in October 2017, 2018, and 2019 and August 2019. The water temperature at each of the times of sampling was consistently 14-15°C. The upstream source water runs approximately 1 mile from its spring source, receives ample natural sunlight, and possesses a population of wild rainbow trout and other fish species (Supplementary Figure 1). The water from this area is diverted through an enclosed pipe that flows ∼0.5 miles directly into the hatchery building and outdoor raceways. The outdoor raceways remain uncovered and receive abundant sunlight. The hatchery building is entirely covered (minimal natural light) and contains 20 individual epoxy-painted concrete raceways into which water is piped before flowing out of the building and into the river. Hatchery raceway surfaces are regularly scrubbed with a coarse brush and visible debris are removed. Between production lots, hatchery raceways are drained, pressure washed, soaked in a bleach solution (200 parts per thousand), and then allowed to dry. The available water quality parameters of the influent are as follows; 2017: 0.018 mg/L total phosphorus (TP) and 2.116 mg/L nitrate/nitrite. The available water quality parameters of the effluent are as follows; 2017: 0.092 mg/L TP, 2.083 mg/L nitrate/nitrite, 7.2 mg/L dissolved oxygen (DO), < 31 mg/L chloride, and 1.0 mg/L ammonia (pH 8); 2018:– 0.156 mg/L TP, 7.4 mg/L DO, < 31 mg/L chloride, and 0.5 mg/L ammonia (pH 8.1); 2019: 0.074 mg/L TP, 7.8 mg/L DO, < 31 mg/L chloride, and 0.5 mg/L ammonia (pH 7.3).

### Surface Samples

Surface samples were collected using sterile Whatman OmniSwabs (Maidstone, United Kingdom; cat. # WHAWB100035). In the indoor hatchery and outdoor raceways, an approximately 1-m long strip of the target surface was swabbed until the swab was visibly discolored. For the hatchery, polycarbonate baffles suspended in the center of the raceway were swabbed on both the front (flow facing) and back sides at the air/water interface and at 1 swab length below the surface (15 cm) (Supplementary Figure 2). The metal tailscreens were only swabbed on the front side at the air/water interface. The epoxy-coated concrete walls of the raceways were typically swabbed at four depths, including the air/water interface, 1 swab length below the surface (15 cm), 2 swab lengths below the surface (30 cm), and at the bottom portion of the wall adjacent to the floor (variable depth as water levels change during production cycle). The walls along the length of the raceway were typically swabbed towards the front third, directly preceding the location of the first baffle. In 2019, a second location of the wall was also sometimes swabbed toward the back third of the raceway, directly behind the final baffle. For the outdoor raceways, swab samples were only collected from the air/water interfaces of walls. For BCS, wall swabs refer to any surfaces that were swabbed (typically rocks) below the water line at various depths. All swabs were immediately placed on ice and then frozen at -20°C for subsequent shipment and storage.

### Water Samples

Six liters of water were collected for each water sample using sterile Whirlpak bags (Madison, WI, USA; cat. # B01447WA). In the hatchery, inflowing and outflowing water samples were typically collected at the inflow and outflow pipes of individual raceways. A few samples were collected from the “global” inflow, which refers to water collected from the primary water pipe before being diverted to individual raceways. Global outflow samples were collected from the main water pipe leading from the hatchery to the river. All water samples were filtered in 1-L aliquots using a previously described system and protocol (22). A 2 µm pore size prefilter (Sigma Aldrich, St. Louis, MO, USA; cat. # AP2502500) and a 0.2 µm pore size Sterivex filter unit (Fisher Scientific, Waltham, MA, USA; cat. # SVGPL10RC) were collected. Attempts to filter the water with a Sterivex filter unit alone (to attain a single representative water sample) proved unfeasible, as clogging was a common occurrence. Prefilters and Sterivex units were immediately frozen at -20°C for shipment and storage.

### Fish Samples

Fish from the outdoor raceways were captured via a net. Gill swabs were collected by direct swabbing of the outward facing gill tissue. Intestinal samples were collected following dissection of euthanized adult fish. Sampling of farmed and wild fish were approved under IACUC Animal Protocol A16-040.

### Diagnosis of CD and BCWD

CD and BCWD were diagnosed by assessing the presence or absence of rod-shaped, oxidase-positive bacteria on gills and within the spleen and kidney tissues of multiple sampled fish.

### DNA Extraction

DNA was isolated from the swab and tissue samples using a Qiagen PowerSoil DNA Isolation kit (Qiagen, Hilden, Germany; cat. # 12888-100) following the manufacturer’s protocol (23). For the prefilter samples, DNA was isolated using a Qiagen PowerWater DNA Isolation kit (cat. # 14900-50-NF). DNA was extracted and isolated from Sterivex filters using the Qiagen Sterivex PowerWater DNA Isolation kit (cat. # 14600-50-NF) following the manufacturer’s protocol. DNA was eluted in 50 µL of buffer EB (Qiagen) and quantified with a Qubit HS dsDNA Assay kit (Life Technologies, Carlsbad, CA, USA). DNA extracts were stored at -20°C until ready for further processing.

### PCR Amplification

For each sample, the V4 hypervariable region of the 16S rRNA gene was amplified using previously validated primers that contained dual-end adapters for indexing as previously described (22, 23). Briefly, PCR reactions were set-up and performed in triplicate using the 515F forward and 806R reverse rRNA gene V4 primers (24) with Illumina MiSeq adaptors. A Bio-Rad C1000 Touch Thermocycler (Bio-Rad Laboratories Inc., Hercules, CA, USA) was used for PCR amplification with the following parameters: 94°C for 3 min followed by 30 cycles of 94°C for 45 sec, 50°C for 60 sec, and 72°C for 90 sec, with a final incubation at 72°C for 10 min.

### Library Preparation and Sequencing

PCR amplicons were prepared for sequencing as previously described (23). Briefly, reactions were pooled, and product presence and size were verified on a QIAxcel (Qiagen). PCR products were then cleaned using a GeneRead Size Selection kit (Qiagen; cat. # 180514) and diluted to 4 nM for sequencing on an Illumina MiSeq (Illumina, San Diego, CA, USA). A 500 cycle MiSeq Reagent kit v2 (Illumina; cat. # MS-102-2003) was used for paired-end sequencing of all libraries.

### Bioinformatic Processing

16S rRNA V4 gene sequence data were demultiplexed through BaseSpace (basespace.illumina.com), after which paired-end data were downloaded and imported into QIIME 2 (version 2021.4) (25). Then, the reads underwent filtering, denoising, and dereplication using the DADA2 denoise-paired plugin (26). Sequences were taxonomically classified and compiled using the feature-classifier (27) and taxa (https://github.com/qiime2/q2-taxa) plugins. Taxonomy was assigned using a pre-trained Naïve Bayes classifier based on the Silva 132 99% OTU database (https://www.arb-silva.de/documentation/release-138/) (28), with the reference trimmed down to the region bound by the 515F/806R primer pair. The default QIIME 2 processing parameters were used throughout the workflow except where otherwise noted.

ASV counts, taxonomy information, and a phylogeny were imported into RStudio (version 1.1.447) (29) from QIIME 2 using the qiime2R package (30). Phyloseq (version 1.36.0) (31), ggplot2 (version 3.3.5) (32), and microViz (version 0.7.9) (33) were used to calculate alpha and beta diversity as well as to generate barplot, boxplot, and ordination visualizations. Statistical testing was performed primarily using the vegan (version 2.5-7) (34) package.

### Controls and QC

DNA extraction controls were processed for every extraction kit lot and used to assess potential kit contaminants and contamination occurring during DNA extraction. PCR negative controls were run with every PCR plate using molecular grade water in place of DNA to assess PCR reagent contaminants and contamination occurring during PCR. Mock community DNA controls (ZymoBIOMICS, Irvine, CA, USA; cat. # D6306) were also amplified during each PCR and used to assess PCR amplification biases.

QC steps performed during library preparation included the previously mentioned use of a QIAxcel to verify product presence and size. For the 2019 samples, 16S-targeted qPCR (35) was performed to verify that the total bacterial load was sufficient to overcome potential well-to-well contamination that may occur in plate-based PCR setups. For this specific group of samples, any sample with more than 10,000 copies of the 16S rRNA gene (as determined by qPCR) was retained (total samples retained, 431/446, 97%). A number of samples did not return reads following sequencing, possibly due to failed PCR reactions (357/431, 83%). Following sequencing and initial processing, only samples with at least 10,000 reads were retained for downstream analyses to guard against the effects of potential contamination on low biomass samples and increase the robustness of downstream analyses (163/357, 46%). Sequence quality filtering and chimeric sequence removal were also performed within QIIME 2 using the default parameters. Five separate MiSeq sequencing runs yielded 17,226,236 paired reads, and after the aforementioned QC steps were 6,131,752 with an average of 46,103 reads and a median of 39,325 reads per sample. The minimum and maximum read counts were 10,720 and 172,746, respectively. A total of 25,790 ASVs were recovered across all samples.

### Statistical Analysis

Alpha and beta diversity index calculations were performed on data rarefied to 10,000 reads. PERMANOVA testing was used to determine statistical significance of group differences for beta diversity indices. Shapiro-Wilk normality testing was used to justify specific statistical testing choices. Normally and non-normally distributed data were analyzed with t-tests and Wilcoxon signed-rank tests, respectively. Holm corrections were used to correct for testing with multiple comparisons, where appropriate. ANCOM-BC default parameters were used, and as recommended, the structural zero flag (neg_lb) was set to “True” only for comparisons in which both groups had 30 or more samples per group. Tested classes had to be present in 10% of samples to be included. For all tests, a corrected p-value of 0.05 was required to achieve significance.

## Results

### Sample Summary

A total of 448 samples were collected at an aquaculture facility between 2017-2019, 163 of which were analyzed using an amplicon sequencing approach. As such, this study represents one of the most extensive microbial surveys conducted on an aquaculture facility and provides important insight into microbial community composition across various sites at the farm, including but not limited to those colonizing the surfaces of raceway walls and other raceway structures, those residing within the water column, as well as those of fish skin and gills. The large and diverse dataset provides an opportunity to compare and correlate microbial communities according to specific farm site (hatchery, outdoor raceway, source water location, inflow, outflow, etc.), sample type (water, surface, etc.), and clinically diseased versus healthy fish populations.

### Microbial Communities of Box Canyon Spring, the Hatchery, and the Outdoor Raceways

We first compared the three sampling sites to determine whether the surface or water communities at each location were distinct from one another and assess if the type of sample or year of sampling were correlated with microbial community structure. The microbial communities of BCS (n = 15), the indoor hatchery facility (n = 134), and the outdoor raceways (n = 14) were compared, with water and surface biofilm samples sampled at all three sites. Additionally, intestine and gill swab samples were included from fish contained in the outdoor raceways to assess the degree of relatedness between host-associated and host-independent samples. Weighted Unifrac distances were calculated for all samples, and PCoA plots were prepared (Figure 1). Microbial community composition was largely dependent on sample type and sampling site. Sterivex and prefilter samples clustered separately from the surface (baffle, wall, tailscreen) samples (Figure 1A) indicating a different overall community composition in biofilms. Fish-associated samples (gills and intestines) clustered closer to the water samples (prefilters and Sterivex). Given the observed differences in microbial community composition between the surfaces (walls, baffles, and tailscreens) and the water (prefilter and Sterivex), these sample types were plotted and analyzed independently. Figure 1B only contains the surface samples (excludes water and fish samples) and shows distinct surface community clustering based on sampling site with BCS and the indoor hatchery resolving into separate clusters, whereas most of the outdoor raceways fell in between these clusters. Figure 1C shows only water samples (prefilters and Sterivex) and lacks obvious clustering by sampling site, whereas a clear separation was noted between the prefilter and Sterivex sample types. This result indicates that the compositions of the communities residing in the water are fairly consistent from the source water to the hatchery and outdoor raceways. Bray-Curtis, Jaccard, and Unweighted Unifrac metrics were also calculated and are displayed in Supplementary Figures 3-5. These results showed overall agreement with those of the weighted Unifrac analysis, with the most obvious divergence being that the hatchery surface samples appeared more variable (larger confidence interval). However, the observed separation between surfaces at different sites and the lack of separation noted for water samples was still evident.

**Figure 1:**
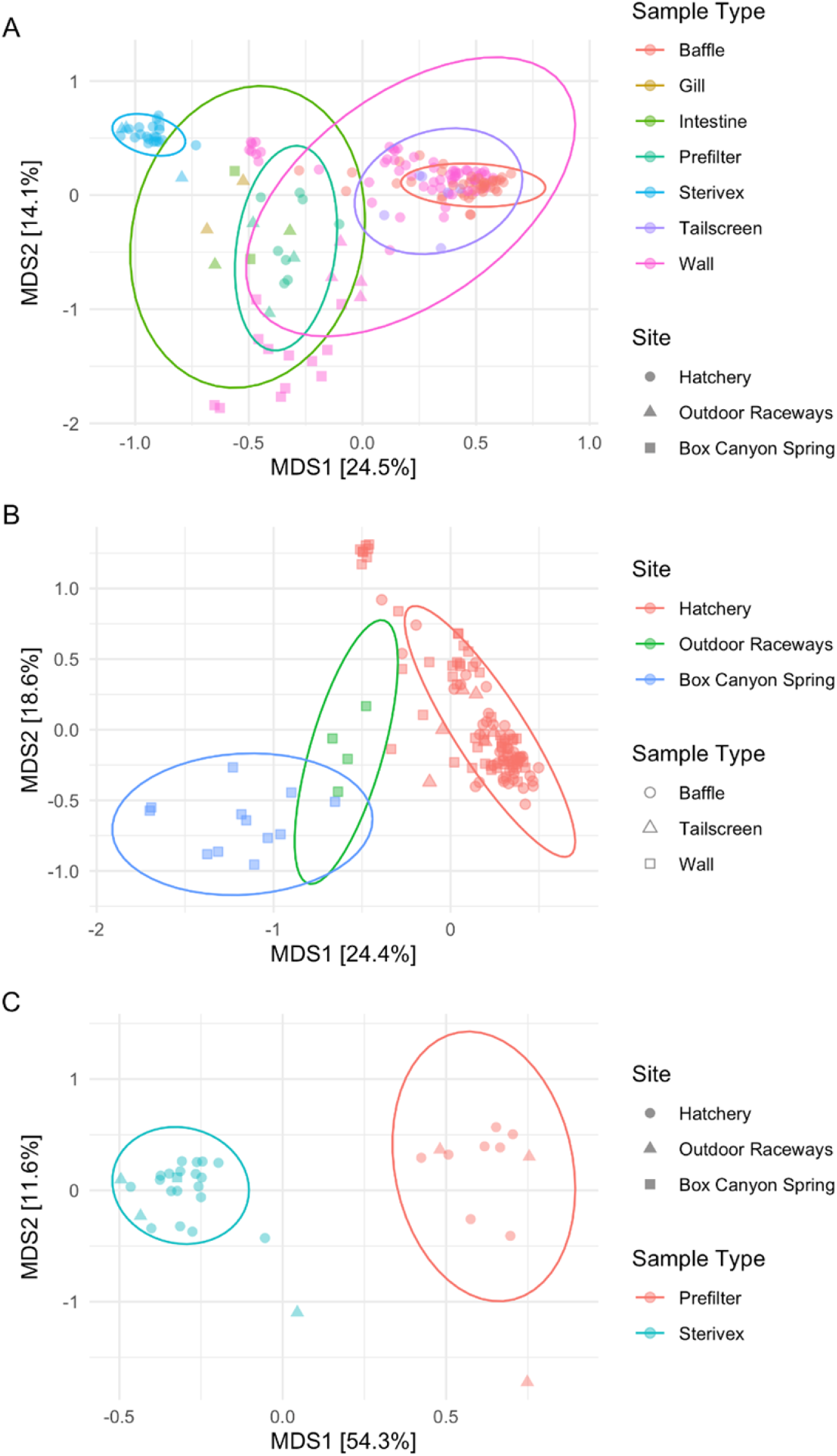
Weighted Unifrac Ordinations of the Three Sampling Sites. (A) All sample types, (B) surface samples only, and (C) water samples only from three sampling sites with 95% confidence ellipses. Three samples or more were required for an ellipse to be drawn for a particular category. Samples are colored by most significant contributing variables.

Next, we tested for the significance of these observed community differences that were attributable to sample site, sample type, and year of sampling (as well as their interactions) using PERMANOVA analysis of surface and water samples (Table 1). For surface biofilm samples, significant differences were observed when testing the effects of sample type, sample site, and year of sampling as well as the interaction of sample type and year and sampling site and year. The largest proportion of variance explained was due to sample site (R^2^ = 0.2048), with all other variables and interactions having R^2^ values <0.10. Significant differences were noted for year but also for the interaction between sampling year and sample site/sample type, indicating that differences due to year of sampling may be due to differences in the types or locations samples were collected from in a given year as opposed to differences solely based on time. Beta dispersion testing was also performed to evaluate differences in dispersion between groups and aid in the interpretation of the PERMANOVA results. Significant differences in dispersion were noted between the walls and baffles (p = 0.0002) and the years 2017 and 2018 (p = 0.003) as well as 2017 and 2019 (p = 0.011). No significant differences were observed between the dispersion levels of sampling sites, further supporting the conclusion that the surface biofilm communities at the three sampled locations are distinct.

**Table 1:**
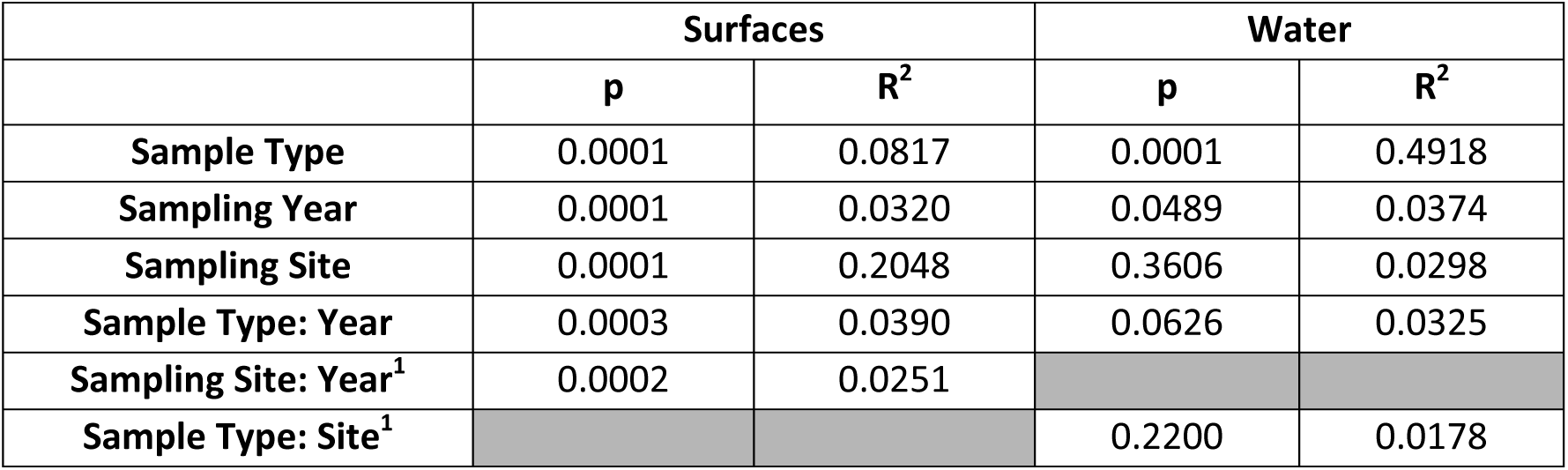
PERMANOVA Testing of Weighted Unifrac Distances for Surface and Water Samples at the Three Sites.

|  | Surfaces |  | Water |  |
| --- | --- | --- | --- | --- |
|  | p | R<sup>2</sup> | p | R<sup>2</sup> |
| Sample Type | 0.0001 | 0.0817 | 0.0001 | 0.4918 |
| Sampling Year | 0.0001 | 0.0320 | 0.0489 | 0.0374 |
| Sampling Site | 0.0001 | 0.2048 | 0.3606 | 0.0298 |
| Sample Type: Year | 0.0003 | 0.0390 | 0.0626 | 0.0325 |
| Sampling Site: Year<sup>1</sup> | 0.0002 | 0.0251 |  |  |
| Sample Type: Site<sup>1</sup> |  |  | 0.2200 | 0.0178 |

<sup>1</sup>Due to sample sparsity for specific sites, testing could

For water samples, sample type (prefilter vs. Sterivex) and sampling year were both significant factors, with sample type explaining a large amount of variance and sampling year explaining a smaller portion (Table 1). No interactions between variables were found to be significant. The beta dispersion results showed significant differences in dispersion between BCS and both the outdoor raceways (p = 0.0014) and hatchery (p = 0.0132). Additionally, dispersion differences were noted between the years 2017 and 2019 (p = 0.0355).

Similar results to the ones found from weighted Unifrac were obtained from testing on Bray-Curtis, Jaccard, and unweighted Unifrac distances as well (Supplementary Tables 2-4). The primary differences between the non-phylogenetic and phylogenetic metrics were smaller R^2^ values contributed by sampling site for surface samples and smaller R^2^ values contributed by sample type for water samples. These results indicate that sampling site has a significant and strong impact on the microbial community structure of the tested surfaces. This contrasts with the water samples where sampling site was not a significant factor influencing the microbial communities. Sample type was the primary significant variable explaining close to 50% of the variance in water samples (particle-associated prefilter vs. planktonic Sterivex). The beta dispersion analysis showed non-significant effects for these aforementioned variables, indicating that uneven variance between groups was not the driver of these significant differences. Significant results were obtained for the sampling year variable for both water and surfaces samples. However, uneven sampling on a yearly basis (significantly more samples in 2019 than 2017 and 2018 as well as only sampling certain sites on a given year) may have been a driving factor for this detected difference and this could be reflected in the significant differences in the beta dispersion results.

Pairwise differential abundance testing using the ANCOM method showed that 36 bacterial classes were present at significantly higher relative abundances within the surface biofilms at the individual sampling sites (Supplementary Table 5). Between the hatchery and outdoor raceways, a significantly higher relative abundance of Cyanobacteria (W = 22.49) was noted in the outdoor raceways. This difference in Cyanobacteria abundance was also observed when testing BCS and the hatchery (W = 14.58) with significantly higher levels found at BCS. This result is likely due to the hatchery receiving minimal natural sunlight, whereas BCS and the outdoor raceways receive ample amounts to support Cyanobacteria and algae. Additionally, significantly higher relative abundances of Deinococci were found at BCS (W = 3.51) and the outdoor raceways (W = 16.61) compared to the hatchery, possibly owing to the survivability of this bacterial class in high UV environments (36). Alphaproteobacteria were detected at significantly higher relative abundances in the hatchery compared to BCS (W = 9.61) and the outdoor raceways (W = 5.06), possibly indicating that this group fills the niche of photosynthetic microbes in low light environments and/or that high densities of fish support and seed Alphaproteobacteria.

### Microbial Communities of the Indoor Hatchery Structure, Biofilms and Water

Three primary structural components of each hatchery raceway [the epoxy-coated concrete walls of the raceways (n = 57), the plastic baffles (n = 43), and the metal tailscreens (n = 7)] were extensively sampled, and the associated microbial communities were analyzed. The inflowing and outflowing water was also filtered, and the microbial communities were profiled in two fractions [a 2 µm or larger prefilter fraction (n = 8) and a 0.2-2 µm Sterivex fraction (n = 19)]. Since the prefilters comprised larger pore diameters than the Sterivex filters, we expected the microbial community sequenced from there to be more enriched in aggregate-lifestyle or particle-attached microbial cells, while the Sterivex data would show the remaining size-filtered fraction, largely represented by planktonic cells.

The total ASV count amongst these samples following filtering was 19,061. Twenty-five classes belonging to 18 phyla comprised the bulk of the microbial communities along with a group of reads unable to be classified to the phylum level (Bacteria_Kingdom) (Figure 2, Supplementary Figure 6). The most dominant classes (totaling >90% of the community composition of each sample type) were also compared using boxplots (Supplementary Figure 7). The surface samples were predominantly composed of the classes Bacteroidia, Gammaproteobacteria, Alphaproteobacteria, and Verrucomicrobiae (Figure 2). These classes were also major components of the microbial community within the water samples, although at generally lower relative abundances, and additional classes were present that are typically found in water (Figure 2, Supplementary Figure 7).

**Figure 2:**
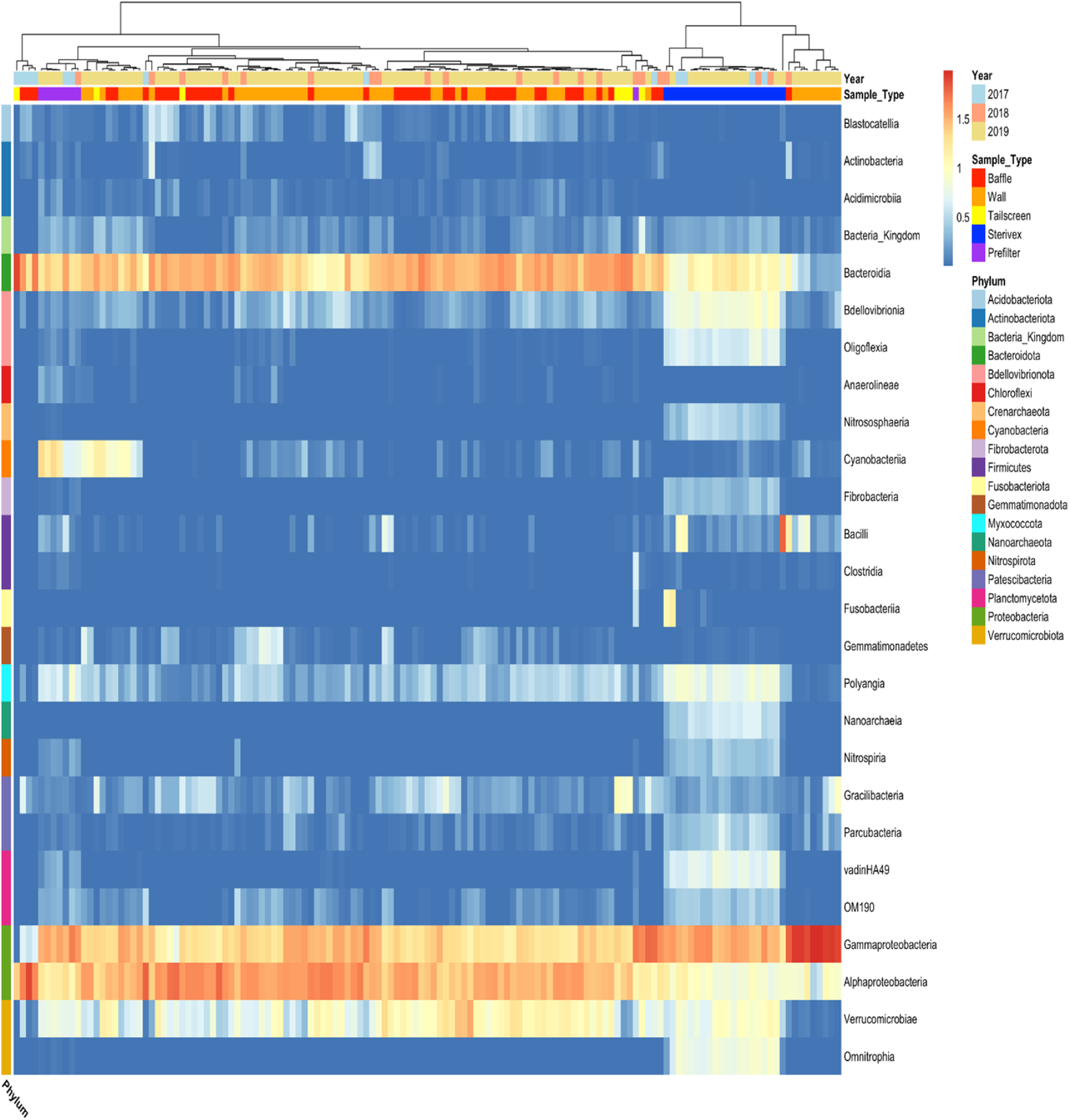
Log-Scaled Heatmap of Bacterial and Archaeal Abundances at the Class Level in the Hatchery. Per sample composition is presented vertically, and samples have been clustered using Pearson correlation values. Phyla are annotated on the left, and classes are grouped based on phylum.

Phylogenetic (weighted and unweighted Unifrac) and non-phylogenetic (Bray-Curtis and Jaccard) beta diversity metrics were used to compare sample types within the hatchery (Figure 3). Sterivex and prefilter samples clustered independently from one another and all other sample types while maintaining a consistent microbial community profile. Surface biofilm samples typically showed greater variation in composition as demonstrated by larger confidence ellipses. Global PERMANOVA testing showed that sample type was a significant variable (Supplementary Table 6). Pairwise PERMANOVA results with multiple testing correction was performed for Bray-Curtis and Jaccard distances and showed significant differences between all sample types (all comparisons, p = 0.01).These results indicate that the communities that reside on the surfaces in the hatchery are compositionally distinct from the seeding water communities, and they are also significantly different from one another.

**Figure 3:**
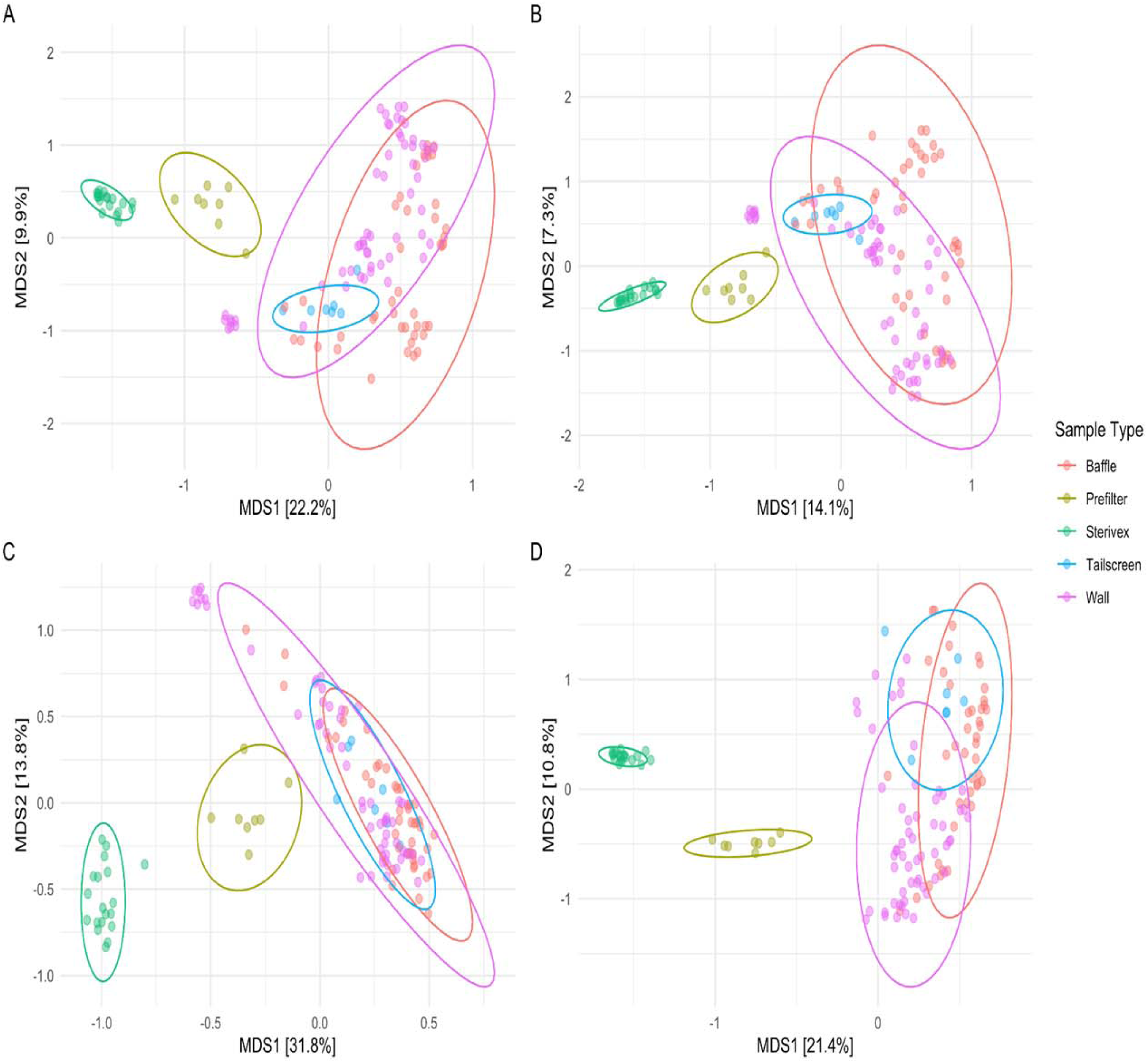
Beta Diversity by Sample Type in the Hatchery. (A) Bray-Curtis, (B) Jaccard, (C) Weighted Unifrac, and (D) Unweighted Unifrac distance metrics colored by sample type with 95% confidence ellipses.

Alpha diversity was calculated for all sample types (Figure 4). The water samples had significantly higher alpha diversity index values compared to the surface samples with the Sterivex samples having the highest. Amongst the surface samples, the walls had the highest Shannon index values followed by the baffles and then the tailscreens. Wilcoxon signed-rank tests were performed in a pairwise fashion for all groups and multiple comparisons correction was applied. For Shannon diversity, all comparisons were shown to have statistical significance (p < 0.05) (Supplementary Table 7). For Faith phylogenetic diversity, all comparisons yielded significant differences (p < 0.05), excluding the comparison between baffles and tailscreens (p = 0.125) and Sterivex and prefilters (p = 0.125) (Supplementary Table 8). These results indicate that the microbial diversity of the water communities was typically greater than that of those residing on the surfaces and that surface type has a significant effect on the community richness and evenness, as the walls had consistently and significantly higher alpha diversity values than baffles or tailscreens.

**Figure 4:**
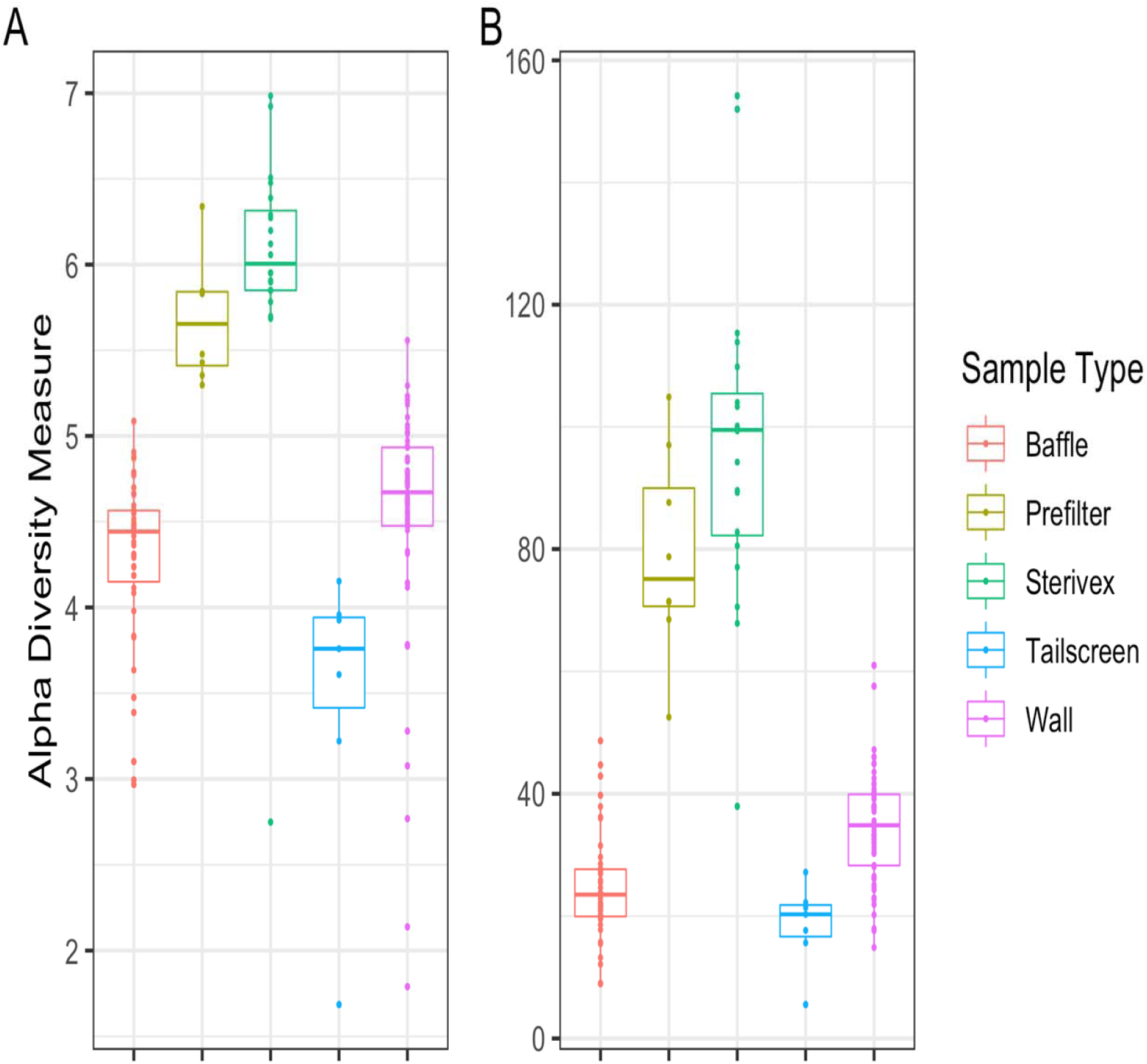
Alpha Diversity by Sample Type in the Hatchery. (A) Shannon diversity index and (B) Faith phylogenetic diversity values were calculated for all samples and arranged by sample type. Samples were rarefied to 10,000 reads before calculating alpha diversity index values.

A presence/absence comparison of the class composition by sample type (only including the inflowing prefilter and Sterivex water samples) showed that most of the microbial classes (13 out of 24) were shared between every sample type (Figure 5A). However, there were 7 classes detected in all samples excluding the tailscreens, 2 classes detected in only the water and on the walls, 1 class detected in only the water and on baffles, and 1 class only detected in water samples. At the family level, 88 of the 211 represented families were present in all sample types (Figure 5B). An additional 28 families were only detected in the water samples, with 8 of those 28 being found exclusively in Sterivex samples. There were also 8 families not observed in Sterivex samples but present in prefilters and surface biofilm samples. There were 57 families detected in all sample types except tailscreen swabs, potentially due to the lower number of tailscreen samples analyzed compared to baffle and wall samples or actual differences in composition. These results reinforce the above alpha diversity results showing that the water communities are far richer than the surface communities. In addition, these findings revealed that 28 microbial families seemingly fail to colonize the surface biofilm communities and that specific surfaces such as the walls contain microbial families (4 present only in water and walls) that baffles and tailscreens do not.

**Figure 5:**
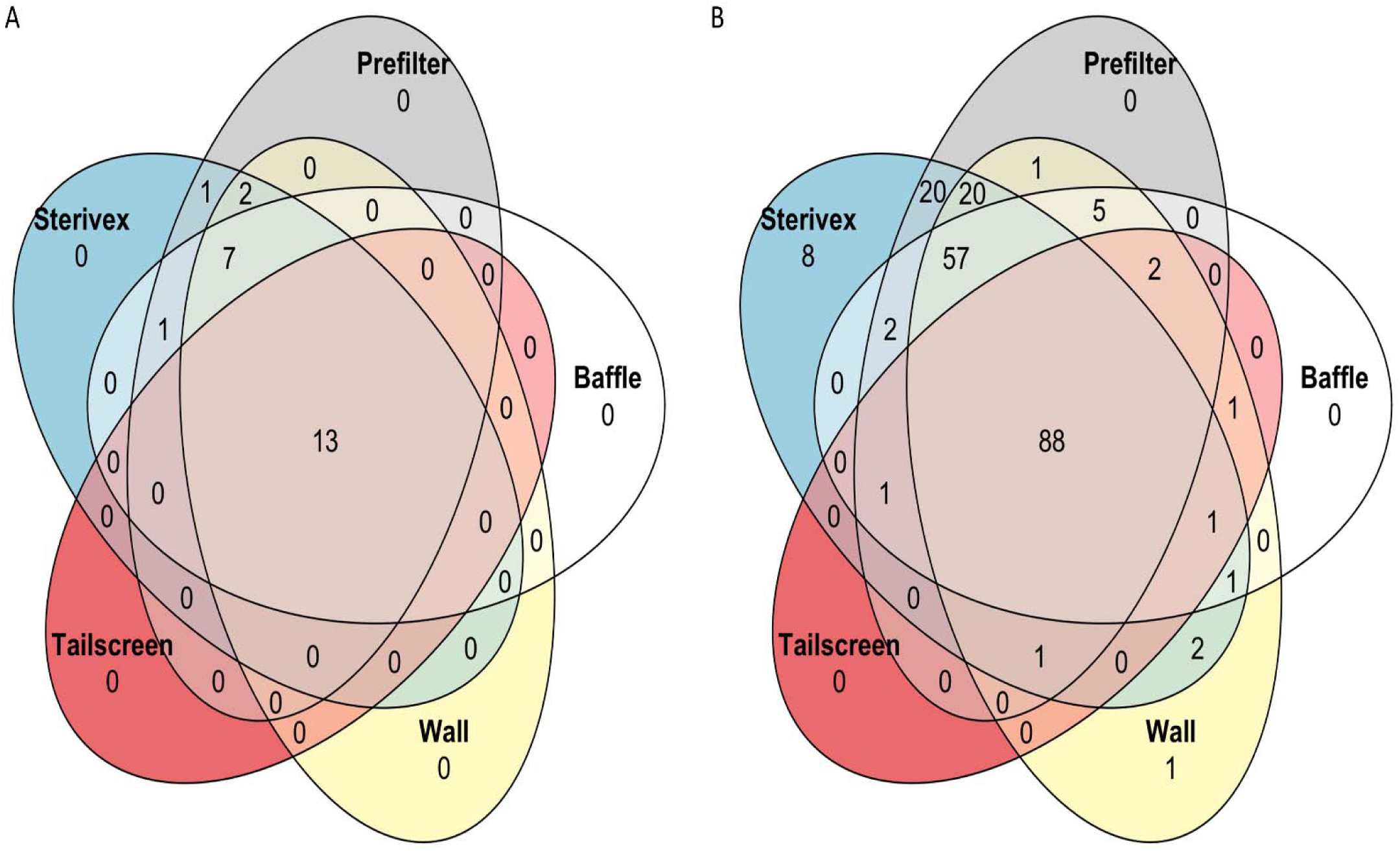
Venn Diagrams of Shared Microbial Classes and Families by Sample Type in the Hatchery. Presence/absence summary of (A) shared microbial classes and (B) shared microbial families stratified by sample type. Microbial classes had to contribute at least 1,000 reads to one or more samples to be included in this comparison. Microbial families had to contribute at least 100 reads to one or more samples to be included in this comparison. Only inflowing water samples were included.

ANCOM-BC testing was used to compare the microbial communities of the three surface types at the class level (Supplementary Figure 8). This comparison showed that 8 bacterial classes were at a significantly higher relative abundance on walls than baffles, 9 were significantly higher on baffles than walls, 12 were significantly higher on walls than tailscreens, 1 was significantly higher on tailscreens than walls, and 11 were significantly higher on baffles than tailscreens. Classes consistently detected at significantly higher levels on walls compared with baffles and tailscreens included OM190, Desulfuromonadia, Anaerolineae, and NB1-j. Classes consistently observed at significantly higher levels on baffles compared with walls and tailscreens included Verrucomicrobiae, Armatimonadia, Fimbriimonadia, and Blastocatellia. No classes were consistently detected at significantly higher levels on tailscreens compared with both walls and baffles (Bacteroidia was found at significantly higher levels on tailscreens than walls but not baffles). These results indicate that not only are there differences in the presence/absence of specific microbial groups but also that many shared groups are present at significantly different relative abundances depending on the surface type profiled.

### Pathogenic Flavobacterium Detection in the Hatchery

*F. columnare* and *F. psychrophilum* species-level detection was performed on the same 16S rRNA dataset described above using a previously validated method (23). A total of 8,854 *F. columnare* reads were detected across 64 samples, and 9 *F. psychrophilum* reads were detected from a single sample, with the relative abundances of *F. columnare* ranging from 0.01 to 6.24%. *F. columnare* was detected for every sampling year and was detected in all sample types. In total, 8/20 Sterivex, 6/8 prefilters, 20/43 baffle swabs, 29/69 wall swabs, and 2/7 tailscreen swabs tested positive for the presence of *F. columnare* (Table 2, Supplementary Table 9).

**Table 2:** Percent Abundance and Frequency of Detection of *F. columnare*.

| Year | Raceway ID | Wall Fc% | Baffle Fc% | Tailscreen Fc% | Inflow Sterivex Fc% | Inflow Prefilter Fc% | Outflow Sterivex Fc% | Outflow Prefilter Fc% | Days from First Feed at Sampling | Daily % Mortality at Sampling | Mortality % at Sampling | Cumulative % Mortality | Columnaris Diagnosed |
| --- | --- | --- | --- | --- | --- | --- | --- | --- | --- | --- | --- | --- | --- |
| 2017 | 6 | N/A | 0 (0/2) | 0.06 (1/1) | N/A | N/A | N/A | 1.29 (1/1) | 71 | 0.017 | 1.93 | 1.97 | - |
|  | 8 | N/A | 0 (0/1) | N/A | N/A | N/A | N/A | N/A | 50 | 0.045 | 1.68 | 1.94 | - |
|  | 14 | N/A | 0 (0/1) | N/A | N/A | N/A | N/A | N/A | 64 | 0.119 | 2.96 | 3.57 | (-) |
|  | 19 | N/A | 0.16 (1/1) | N/A | N/A | N/A | N/A | N/A | 57 | 0.007 | 0.99 | 1.05 | - |
|  | 20 | N/A | 0 (0/1) | N/A | N/A | N/A | N/A | N/A | 57 | 0.009 | 1.20 | 1.27 |  |
|  | Global | N/A | N/A | N/A | 0.02 (2/2) | 0.03 (1/1) | 0.13 (2/2) | N/A | N/A | N/A | N/A | N/A | N/A |
| 2018 | 9 | 0.04 (2/3) | 0.02 (3/4) | 0 (0/1) | 0 (0/1) | 0 (0/1) | 0.02 (1/1) | 0.99 (1/1) | 42 | 0.002 | 8.30 | 9.25 | - |
|  | 11 | 0 (0/3) | 0.01 (1/3) | 0.14 (1/1) | N/A | N/A | N/A | N/A | 71 | 0.004 | 3.61 | 3.65 | - |
|  | 16 | 0 (0/1) | N/A | N/A | N/A | N/A | N/A | N/A | 28 | 0.002 | 0.14 | 0.93 | - |
|  | Global | N/A | N/A | N/A | 0 (0/1) | N/A | N/A | N/A | N/A | N/A | N/A | N/A | N/A |
| 2019 | 1 | 0.05 (7/7) | 0.02 (3/7) | N/A | 0 (0/1) | 0 (0/1) | 0 (0/1) | N/A | 62 | 0.017 | 4.10 | 4.25 | (+) |
|  | 2 | 0.31 (6/6) | 0.15 (7/7) | N/A | 0 (0/1) | 0.08 (1/1) | 4.51 (1/1) | 6.24 (1/1) | 50 | 2.562 | 24.22 | 28.84 | + |
|  | 4 | 0.02 (3/5) | 0.02 (3/6) | 0 (0/1) | 0 (0/1) | N/A | 0 (0/1) | N/A | 64 | 0.006 | 14.26 | 14.42 | (+) |
|  | 8 | 0 (0/3) | 0 (0/3) | 0 (0/3) | N/A | N/A | N/A | N/A | 73 | 0.012 | 1.34 | 1.45 | - |
|  | 9 | 0.01 (1/3) | N/A | N/A | 0 (0/1) | 0.04 (1/1) | 0 (0/1) | N/A | 20 | 0.004 | 0.45 | 3.13 | - |
|  | 10 | 0 (0/4) | N/A | N/A | 0.02 (1/1) | N/A | 0.04 (1/1) | N/A | 7 | 0.000 | 0.00 | 9.59 | (-) |
|  | 11 | 0.01 (1/4) | N/A | N/A | N/A | N/A | N/A | N/A | 6 | 0.000 | 0.00 | 8.47 | - |
|  | 12 | 0.03 (4/4) | 0.02 (2/7) | N/A | N/A | N/A | N/A | N/A | 79 | 0.004 | 45.40 | 45.45 | [-] |
|  | 16 | 0 (0/4) | N/A | N/A | 0 (0/1) | N/A | 0 (0/1) | N/A | 20 | 0.086 | 0.90 | 8.82 | (-) |
|  | 17 | 0 (0/4) | N/A | N/A | N/A | N/A | N/A | N/A | 20 | 0.063 | 0.92 | 6.63 | (-) |
|  | 18 | 0.01 (2/6) | N/A | N/A | N/A | N/A | N/A | N/A | 20 | 0.072 | 0.83 | 12.81 | (-) |

(+) = active columnaris cases but post peak, (-) = no active cases but infection occurred sometime following sampling during production cycle, [-] = no active cases but previous large outbreak had occurred, N/A = not applicable due to samples not being collected or being excluded during processing.

Based on recent taxonomic restructuring, *F. columnare* has been divided into four species: *F. columnare, Flavobacterium davisii*, *Flavobacterium oreochromis*, and *Flavobacterium covae* (12). A 16S rRNA V4 gene sequence comparison was performed between the four type strain sequences and the 7 ASVs within this dataset that were assigned to *F. columnare* based on the Silva database (Supplementary Figure 9). The primary ASV in this dataset, comprising 74% of the recovered *F. columnare* reads, was an exact match to the type strain sequence for *F. columnare* as defined within the new taxonomy. An additional two ASVs comprising 13% of the *F. columnare* reads had a 1 bp difference from the type strain of *F. columnare.* Based on this alignment, confidence in discerning between *F. columnare, F. davisii*, and *F. covae* solely using the V4 region of the 16S rRNA gene is tenuous at best, as each species differs by at most 2 bp.

The co-occurrence of *F. columnare* and the clinical manifestation of columnaris within a given raceway was also examined. Of the 12 raceways where *F. columnare* was detected, 3 had active columnaris infections, 1 had previously experienced a large columnaris outbreak but had no known active cases, 2 had no active cases during sampling and no history of columnaris but experienced an outbreak following sampling, and 6 had no diagnosed columnaris infections during the production cycle. Typically, raceways where no columnaris was detected over the course of the production cycle had lower levels of *F. columnare* compared to raceways where columnaris was identified (Table 2). The relative levels of *F. columnare* in samples in which *F. columnare* reads were recovered (one or more reads) are compared in Figure 6. The levels of *F. columnare* in surface biofilm samples in the 6 healthy raceways was shown to be significantly lower than those detected in the 6 diseased raceways (Wilcoxon, p = 0.0028) (Figure 6A). In contrast, the levels of *F. columnare* in water samples in the 6 healthy raceways were not significantly different than those detected in the 6 diseased raceways (Wilcoxon, p = 0.095) (Figure 6B). This result may be in part due to the difference in sample size for these comparisons, as only 9 water samples were included versus 48 total surface samples (5 water samples were global inflow/outflow samples and were not included as disease status could not be attributed).

**Figure 6:**
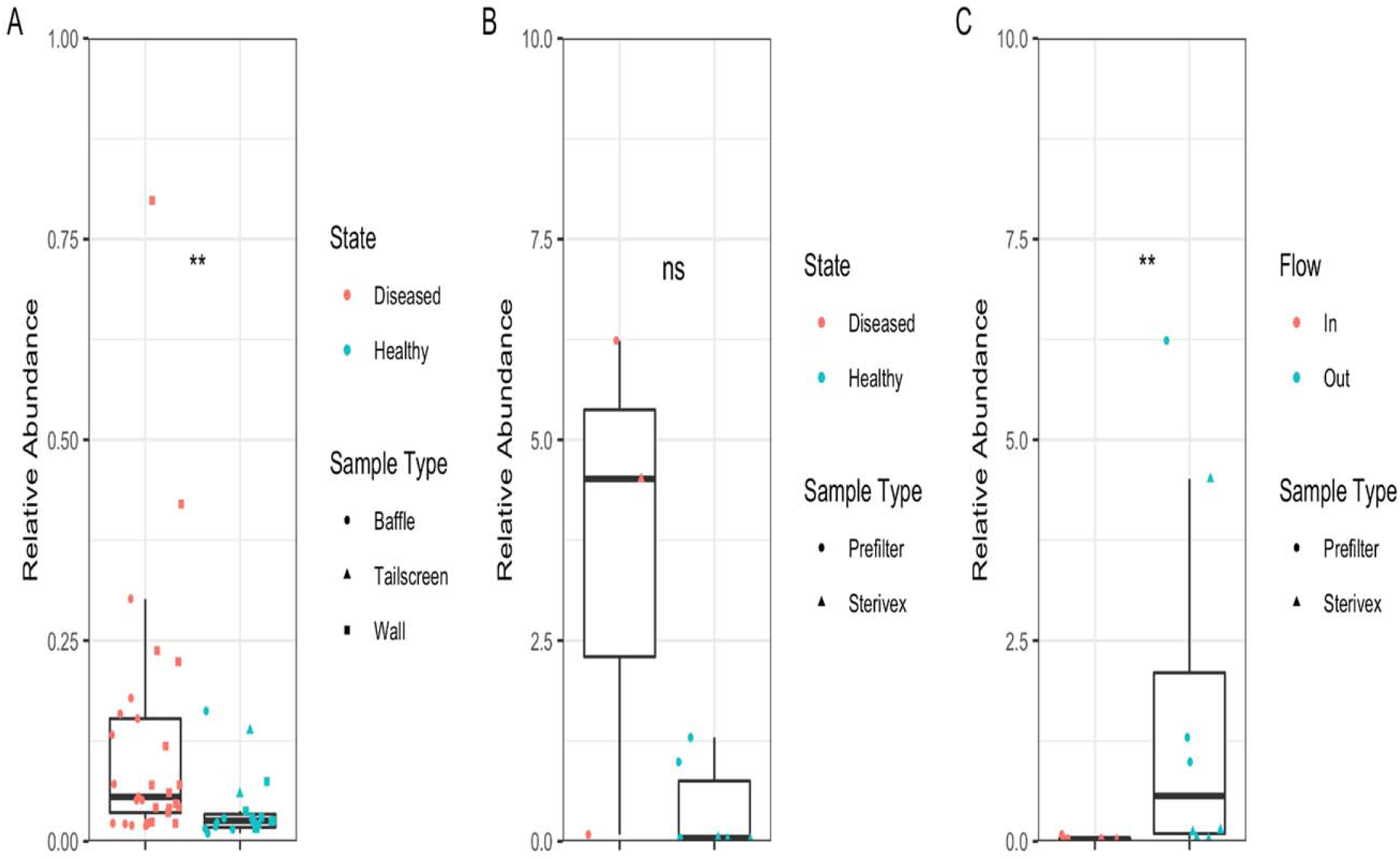
Relative Abundances of *F. columnare* in the Hatchery. Boxplots display the distribution of relative abundances from (A) surface samples grouped by raceway disease status and (B) water samples grouped by raceway disease status and (C) flow direction. Only samples with one or more *F. columnare* reads were included. Global inflow and outflow water samples were not included in disease state figures as water from healthy and diseased raceways would be collected in these samples. Wilcoxon signed-rank tests were performed to test for differences. ns = not significant.

Inflowing and outflowing water samples both contained *F. columnare*, and in multiple cases, *F. columnare* was detected in both the inflow and outflow of the same individual raceway. However, a key difference in these specific cases and with inflow/outflow samples in general was consistently higher levels of *F. columnare* in the outflowing water. In the positive inflow samples, the median percent abundance of *F. columnare* was 0.016 and 0.038% for the Sterivex (n = 3) and prefilter samples (n = 3), respectively. In the positive outflow samples, the median percent abundance of *F. columnare* was 0.114 and 1.294% for the Sterivex (n = 5) and prefilter samples (n = 3), respectively. As these are relative abundances, it is possible *F. columnare* is not truly increasing but that other microbial taxa are decreasing in abundance, causing the percent abundance of *F. columnare* to increase. However, qPCR quantitation of the total microbial load within a subset of 2019 samples showed no significant differences in the total bacterial abundance between inflowing and outflowing prefilter samples (t-test, p = 0.454), indicating that *F. columnare* is likely being amplified in the raceway. Additionally, significantly higher relative abundances of *F. columnare* were detected in the outflowing water samples (n = 8) than in the inflowing water samples (n = 6) (Wilcoxon, p = 0.008; Figure 6C). Furthermore, despite a limited number of samples collected at BCS, *F. columnare* was detected on the piping that flows water to the farm (7 reads) as well as on a submerged rock surface (44 reads). The detection of this pathogen in both the upstream location and the inflowing water reveals BCS as a likely source for recurrent introduction of *F. columnare* into the hatchery and outdoor raceways.

## Discussion

This study focused on the characterization and comparison of the microbial communities associated with the source water, outdoor raceways, and hatchery at a freshwater trout aquaculture facility over three years. The microbial communities of incoming water at inflow, outflow, and within raceways, as well as of surfaces of baffles, tail screens, and raceway walls were characterized. As young fish are particularly vulnerable to *F. columnare* and *F. psychrophilum*, the hatchery building was the most extensively sampled site, and five different sampling location types were characterized, compared, and screened for the presence of these specific flavobacterial pathogens.

The three environments in water-flow succession were compared to assess site-based variation of microbial communities within water and on surfaces. The water source location (BCS), hatchery, and outdoor raceways of the farm had consistent and mostly indistinguishable microbial communities within the water. Sampling site was not shown to be a significant variable contributing to differences in microbial community structure in either the prefilter or Sterivex water fractions. This result indicates that the overall composition of the water community from the natural source spring through to the outflow from the farm is stable and not significantly different despite variations in light availability, animal abundance, and water quality parameters between the three sampled sites. Additionally, sampling year was noted as a significant variable influencing microbial community structure for water samples while only contributing a small amount of variance. Considering the uneven site sampling of water between years (e.g., the outdoor raceways were only sampled in 2017 while the hatchery was sampled 2017-2019), we consider the microbial communities in the water to be consistent on a yearly basis, providing the opportunity for greater reproducibility and stronger study conclusions. As the vast majority of samples were collected during October of each year (a small subset was from August 2019), seasonal variance in community composition may exist that was not explored in the present study.

Compared to the water samples, the surface biofilm communities were significantly different between the three sites. Hatchery surfaces contain a distinct community that differs from the communities residing within BCS and the outdoor raceways. A determining factor in this difference may well be that the hatchery raceways are a poor environment for bacterial (Cyanobacteria) and eukaryotic phototrophs (algae), whereas BCS and the outdoor raceways receive plentiful sunlight. ANCOM results indicated that Cyanobacteria were present at significantly higher levels on BCS surfaces and the outdoor raceway surfaces compared to hatchery surfaces.

Focusing on the hatchery environment specifically, water and surface communities were shown to be highly distinct. Alpha diversity calculations showed that the water communities were significantly more diverse than surface biofilm communities. Additionally, beta diversity results exhibited significantly different clustering of prefilter and Sterivex samples from wall, baffle, and tailscreen biofilm samples. Presence/absence comparison plotting showed every microbial class detected within the hatchery environment was present within the inflowing water community, and all but two microbial families were detected in the inflowing water. This result indicates that while most biofilm constituents are likely seeded directly from the water community, only a distinct subset of this water community can establish itself within the surface biofilm community. It is also worth noting that the potential microbial contributions from the feed as well as the fish themselves was not accounted for in the present study. The two microbial families not detected in the water (Streptococcaceae and Nakamurellaceae) but detected on the surfaces may have originated from one of these two sources. Interestingly, Streptococcaceae were found at moderate levels in fish-associated samples from the outdoor raceways and were observed in outflowing water samples (data not shown), possibly implicating the farmed species as the source of these bacteria. Alternatively, it is possible these microbial classes are present in the water at low enough levels where detection through sequencing is not possible and amplification within the surface biofilms then allows for detection.

Specific surface types within the hatchery were also unique from one another. Walls were shown to have significantly higher alpha diversity than baffles and tailscreens. All three surface types were also shown to cluster in a significantly different manner regardless of the beta diversity metric used. The vast majority of the microbial classes and families detected were present in all three surface type communities but at different relative abundances. This result indicates that despite having identical sources (the water, fish, and feed), specific microbial taxa favor unique surfaces based on potential factors such as material type, surface roughness, nutrient concentrations, hydrodynamics, or other factors(37). Hydrodynamics, surface materials, and exposure time may well explain why the wall, a rough concrete surface located on the periphery of the raceway, has a greater community richness compared to corrugated polycarbonate baffles and stainless steel tailscreens (smoother surfaces located perpendicular to the flow of water(38)). In addition, the baffles are placed into the raceways several weeks after the raceway is seeded with fish, and the water level increases as the fish mature. Microbes with less adept adhesion mechanisms may be able to withstand this reduced flow force along the walls compared to the medial flow force experienced on baffles and tailscreens. There was also an interesting subset of wall swabs that generally clustered distinctly from other wall swabs and surfaces (Figure 3). After reviewing available farm metadata, we are currently investigating the potential for fish age or water contact time as contributors to microbial community composition that may explain this unique clustering pattern.

The lone hatchery sample in which *F. psychrophilum* was detected was a baffle swab from a raceway towards the end of a columnaris outbreak. The detection of *F. psychrophilum* in only this sample may indicate that a coinfection was occurring within this raceway, but diagnosis was not undertaken as an infection cause was already established. A previous study at this farm site (but not at the hatchery facility) identified *F. psychrophilum* on sampled fish at a high frequency using qPCR (39). The lack of *F. psychrophilum*-positive samples in our dataset may be at least partly due to the reduced sensitivity of MiSeq sequencing for pathogen detection compared to traditional qPCR methods (23). *F. psychrophilum* may in fact be present but is simply at too low of a level to be reliably detected using this method.

The sampling of inflowing water shows a clear source of *F. columnare* that can act to reintroduce this and other pathogens from the untreated inflowing water into cleaned and disinfected raceways. *F. columnare* was detected over multiple years in both prefilter and Sterivex samples for inflowing source water. *F. columnare* was also detected within two surface biofilms in the upstream source environment, establishing its presence there. The populations of resident trout and other fish species in BCS together with the ability of *F. columnare* to survive independent of a host provides a reasonable scenario for reintroduction into the hatchery via the inflow. This reinforces the idea that this and other pathogens are likely endemic and the continued use of any approved and available vaccines, even when disease is not present, should be continued to avoid future outbreaks. We also observed that outflowing water samples from specific raceways where *F. columnare* was detected in surface biofilms showed higher relative abundances of *F. columnare* than the inflow. This result indicates the possible amplification of this pathogen in surface biofilms within the raceway followed by shedding into the water where it is then detected in the outflow. Higher levels of *F. columnare* in biofilms and high levels in outflowing water was observed in raceways with no clinical signs of columnaris This finding supports the idea that this shedding likely occurs from either infected, subclinical fish (more fish-associated samples would be needed to investigate this) or from the surface biofilms themselves.

*F. columnare* was detected in all 5 sample types, including, most interestingly, within surface biofilms in both infected and healthy raceways. Intriguingly, *F. columnare* was detected in two swabbed locations of a raceway that had been emptied and mostly dried (data not shown). This result further solidifies the idea that *F. columnare* is ubiquitous in the environment and is adequately equipped with molecular mechanisms to reside within a complex biofilm community. This finding also highlights that the mere presence of these opportunistic pathogens does not necessarily have a direct correlation with a disease phenotype in the farmed species. Fish immune factors and general stress within the farm environment likely predispose fish to infection by opportunistic pathogens residing in the surrounding communities. However, this phenomenon does not completely invalidate the idea that preventing potentially pathogenic microbes from residing within the hatchery environment would be beneficial, as bacterial load likely plays a crucial role in disease occurrence as well. Increased exposure events due to higher levels of *F. columnare* entering the water from surface biofilm sources may also increase the likelihood for infections to occur. Our results showed that infected raceways had significantly higher levels of *F. columnare* in surface biofilms than healthy ones, but more work will be needed to establish whether this is a precursor to infection or the consequence of one.

In the present study, the communities residing on surfaces within a rainbow trout aquaculture facility were shown to be distinct from the water community that initially seeds these communities and are distinct from one another. The presence of the causative agent of a recurrent disease within the facility was also shown to be widespread, with inflowing water and surface biofilms acting as source and sink for *F. columnare*. These results provide valuable insight into a mostly uncharacterized setting and provides a backdrop for future studies concerning microbial ecology of the built environment, biofilm maturation, antibiotic resistance, probiotic strain sourcing, and disease prevention.

## Data Availability

Raw read data are available in the NCBI SRA database under project ID PRJNA821905.

## Code Availability

Code is available at https://github.com/joerggraflab/Code-for-Testerman-Beka-2022.

## Acknowledgements

We thank the University of Connecticut Microbial Analysis, Resources, and Services (MARS) facility for performing the sequencing work. We thank Chris Jackson and Dr. Jesse Trushenski for providing farm metadata. We thank Dr. Jeremiah Marden for providing feedback on the manuscript.

---

**Supplementary Figure 1:**
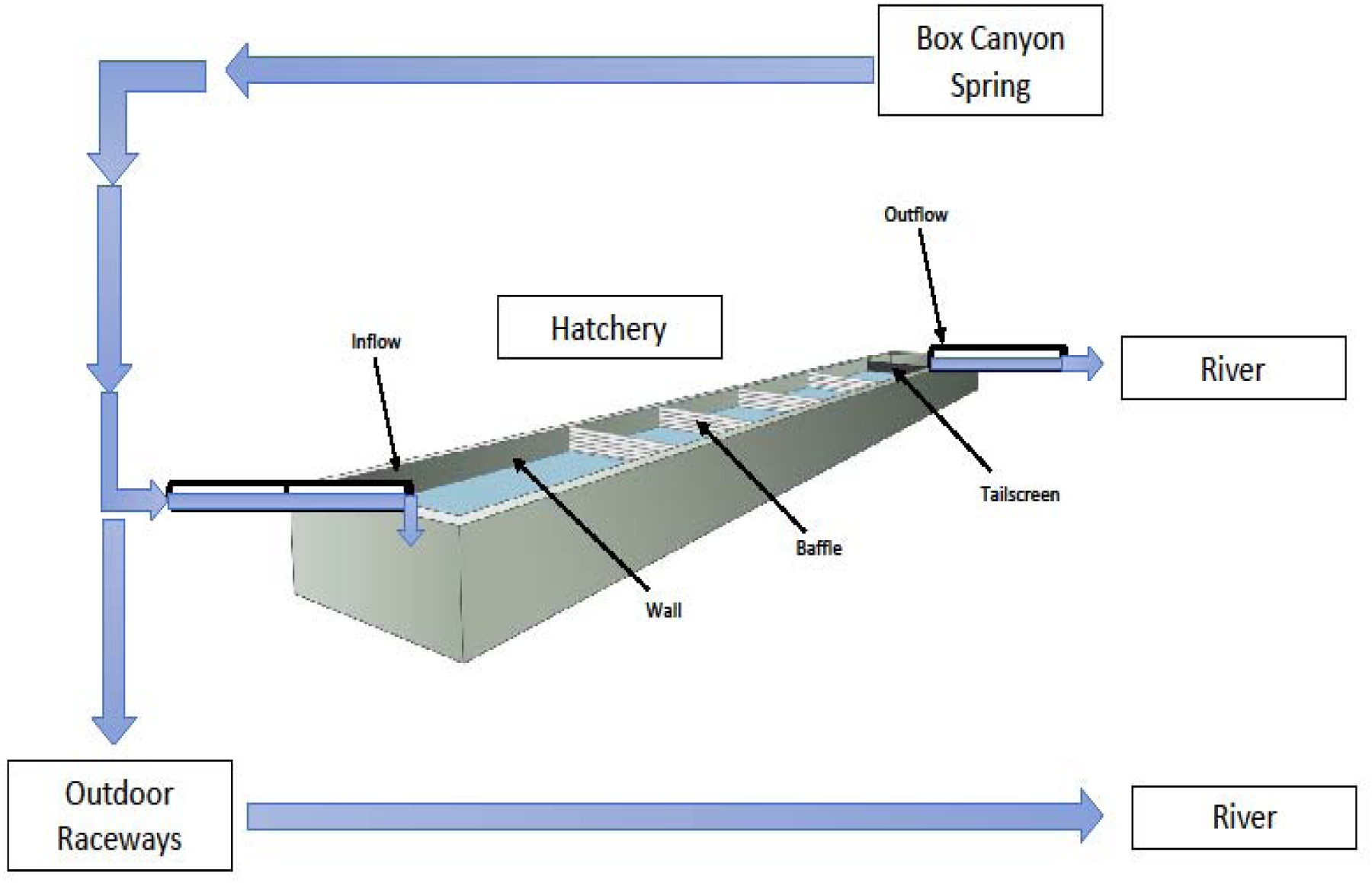
Diagram of Water Flow Progression and Hatchery Raceway Structure. Blue arrows indicate the flow of water starting from the upstream, Box Canyon Spring and proceeding through the hatchery and outdoor raceways before flowing out into the river. This is a simplified layout and is only meant to show the basic progression of water as it pertains to this study.

**Supplementary Figure 2:**
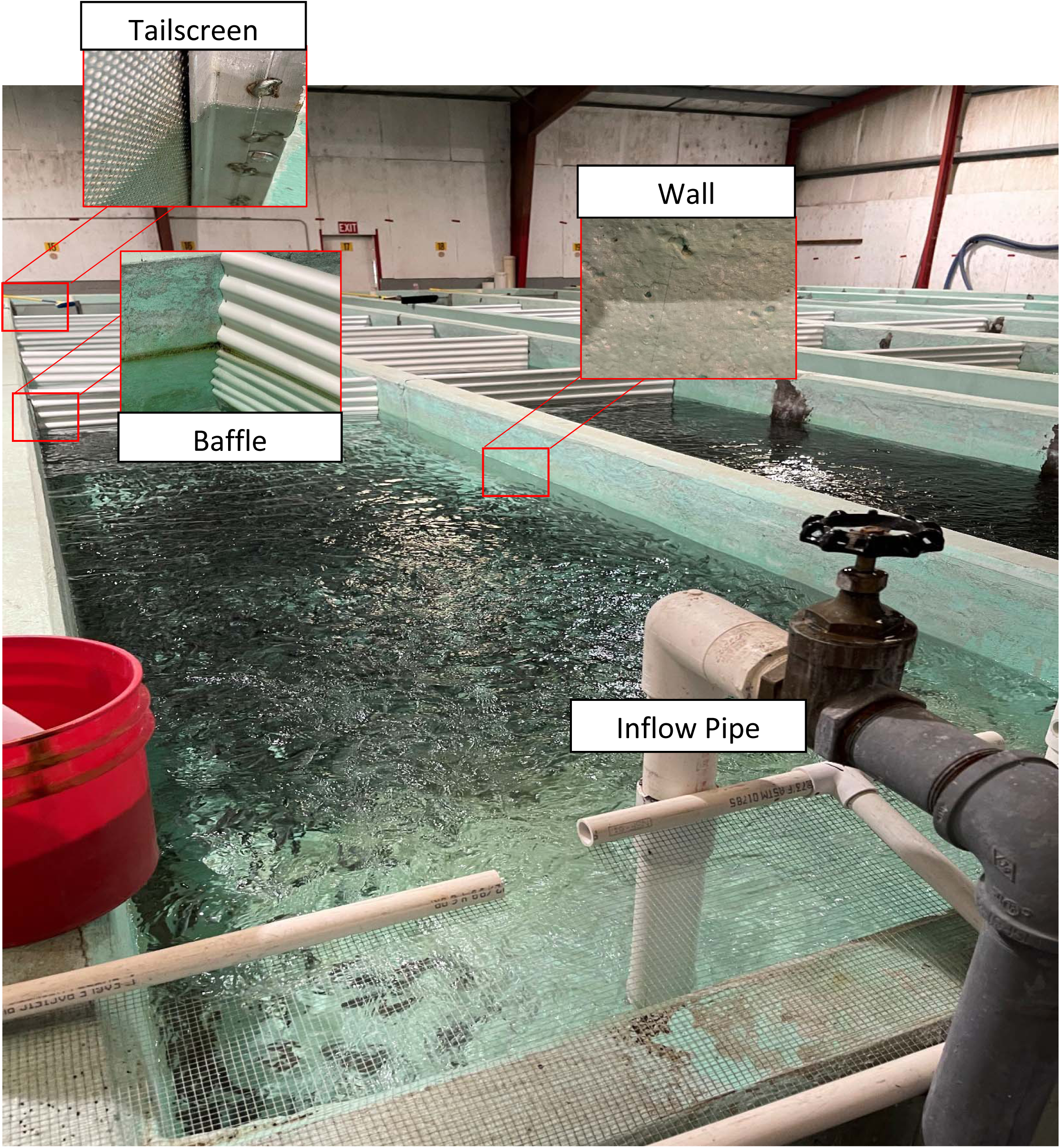
Raceway Layout and Structures. The raceway layout is displayed with the inflowing water pipe and three sampled surface types (wall, baffle, and tailscreen) highlighted. Water flows away from the inflow pipe towards the end of the raceway, where it flows out through a pipe (not shown).

**Supplementary Figure 3:**
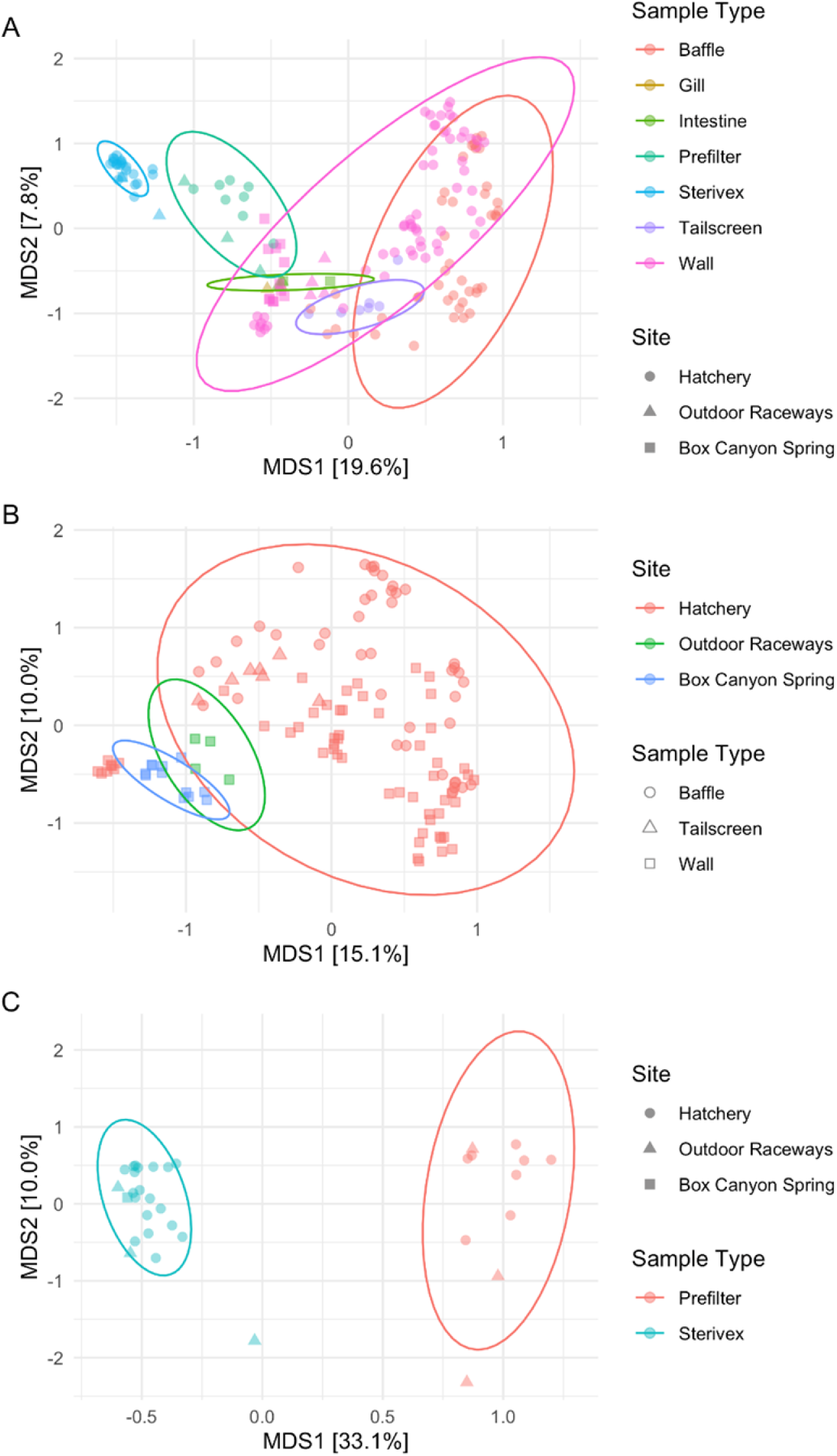
Bray-Curtis Ordinations of the Three Sampling Sites. (A) All sample types, (B) surface samples only, and (C) water samples only from three sampling sites with 95% confidence ellipses. Three samples or more were required for an ellipse to be drawn for a particular category. Samples are colored by most significant contributing variables.

**Supplementary Figure 4:**
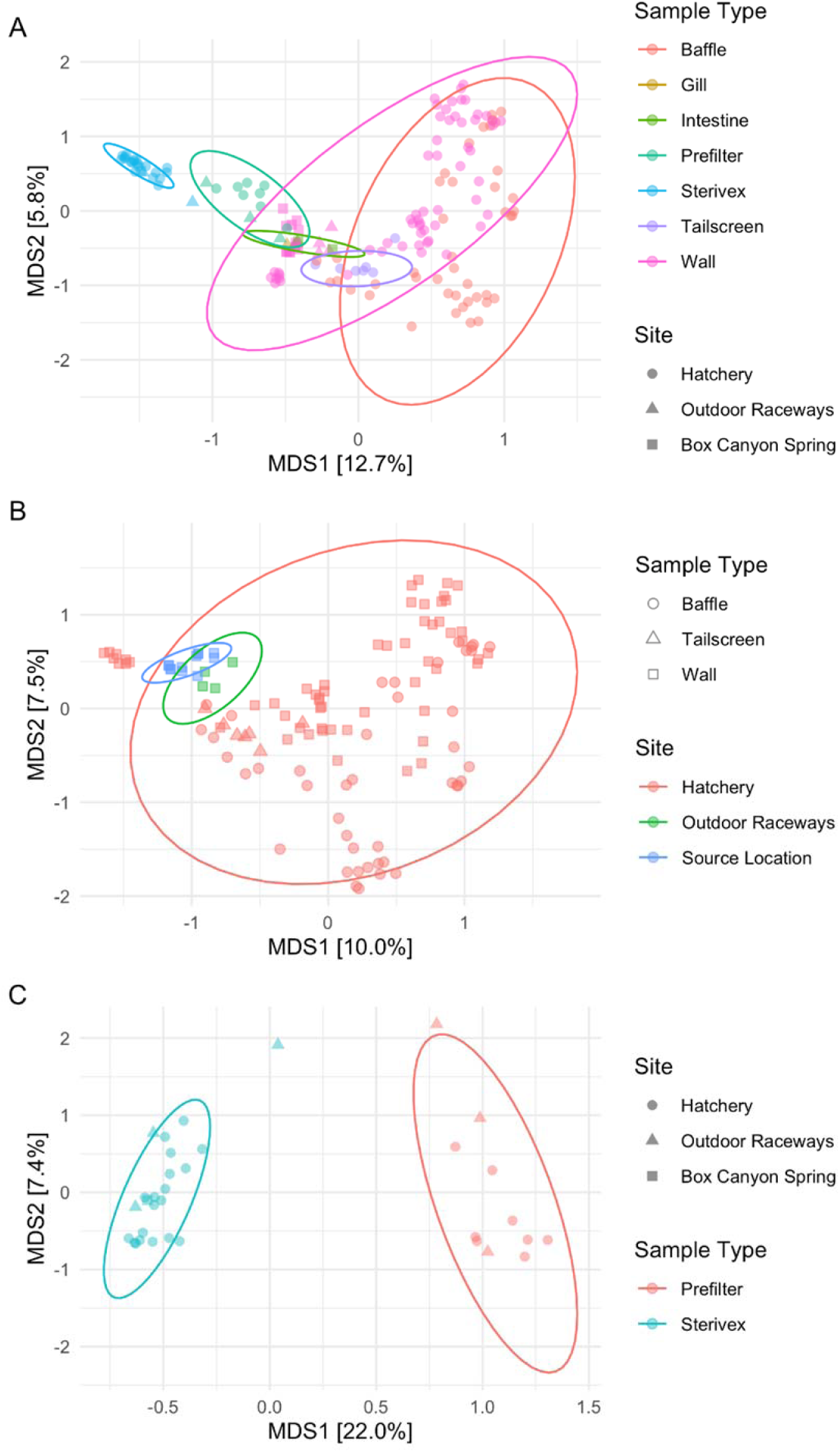
Jaccard Ordinations of the Three Sampling Sites. (A) All sample types, (B) surface samples only, and (C) water samples only from three sampling sites with 95% confidence ellipses. Three samples or more were required for an ellipse to be drawn for a particular category. Samples are colored by most significant contributing variables.

**Supplementary Figure 5:**
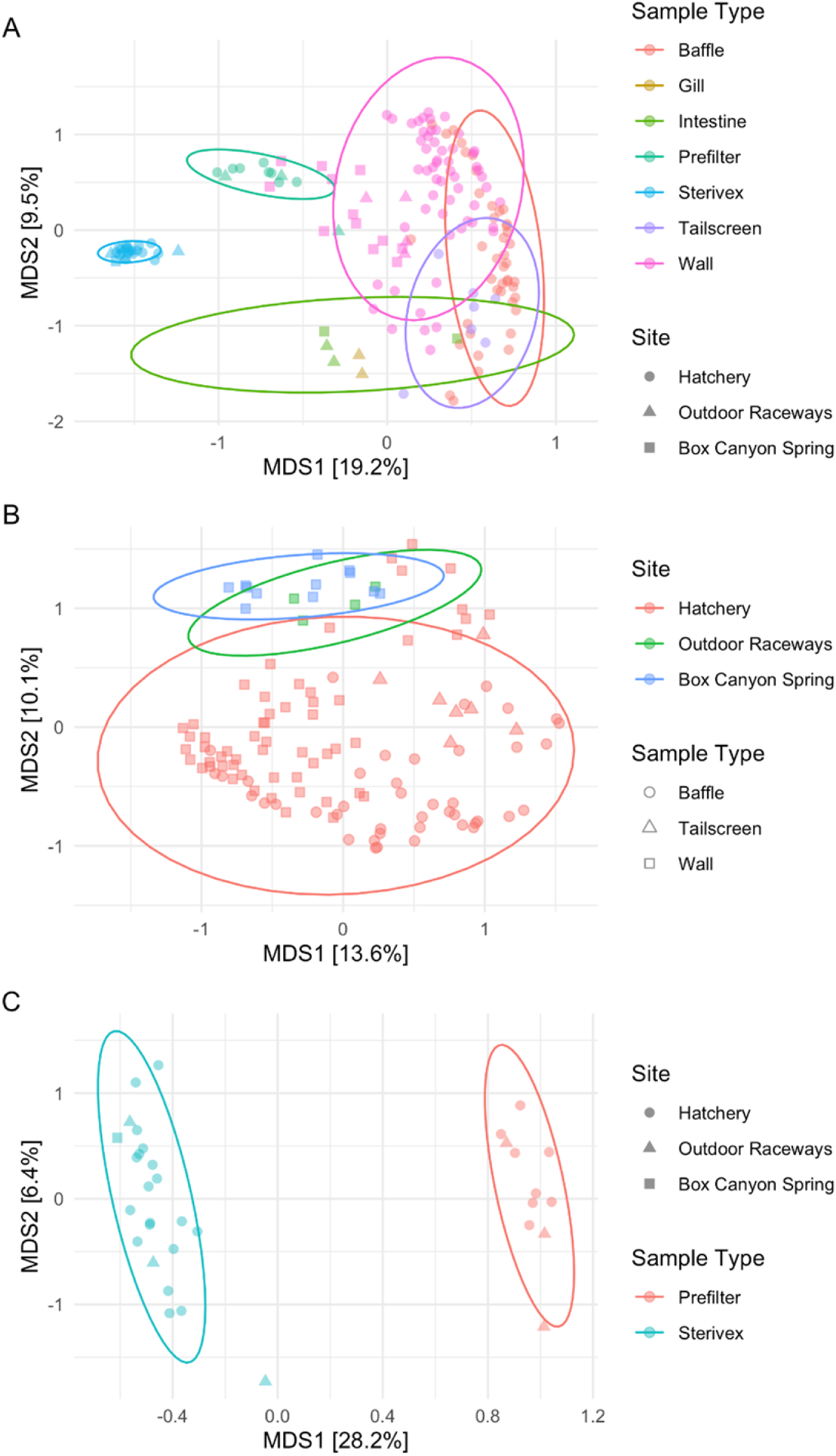
Unweighted Unifrac Ordinations of the Three Sampling Sites. (A) All sample types, (B) surface samples only, and (C) water samples only from three sampling sites with 95% confidence ellipses. Three samples or more were required for an ellipse to be drawn for a particular category. Samples are colored by most significant contributing variables.

**Supplementary Figure 6:**
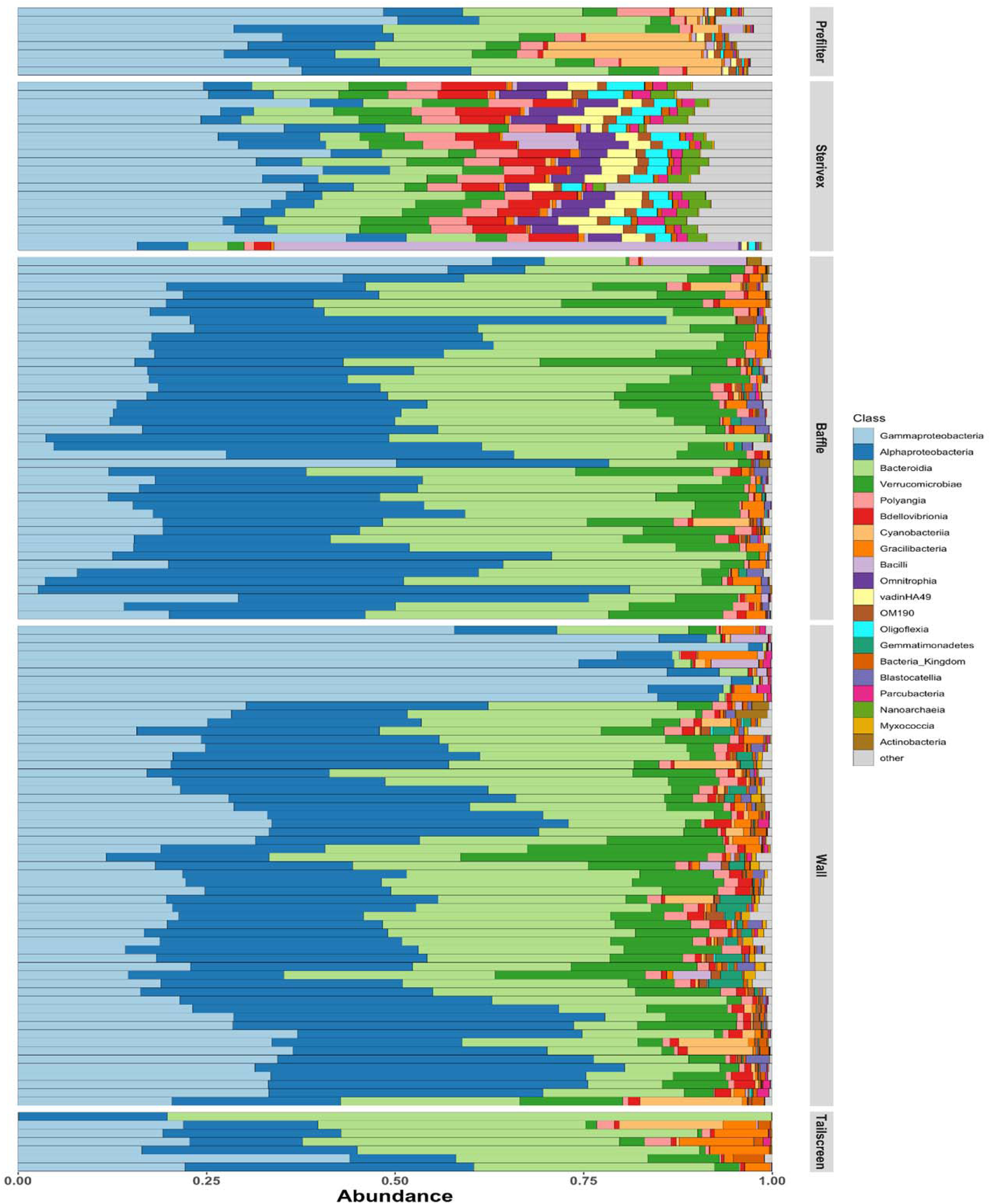
**Class Level Relative Abundance Bar Plots Grouped by Sample Type.**

**Supplementary Figure 7:**
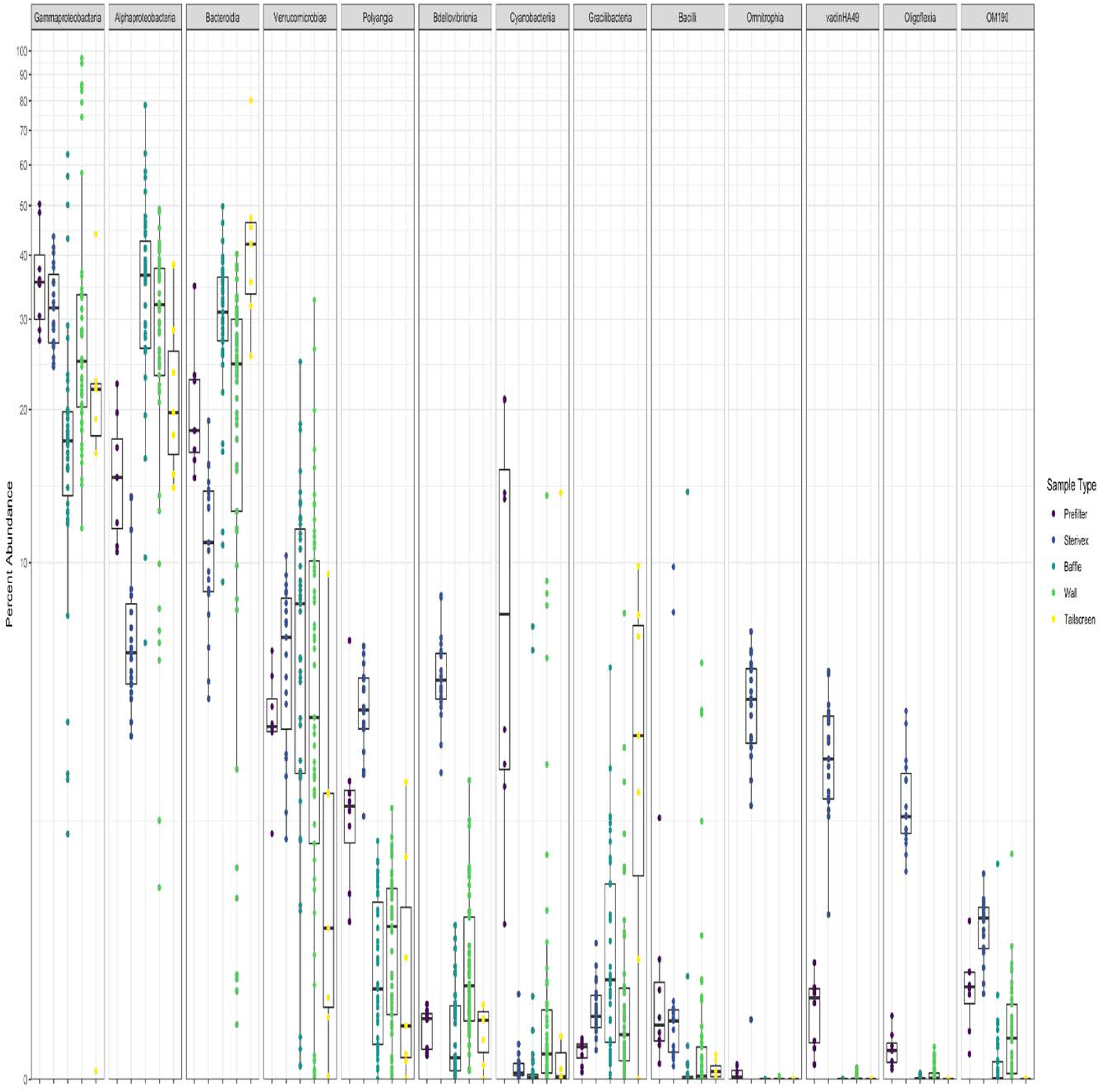
Boxplots of Dominant Bacterial Classes by Sample Type. Percent abundances are shown on the y-axis on a log scale.

**Supplementary Figure 8:**
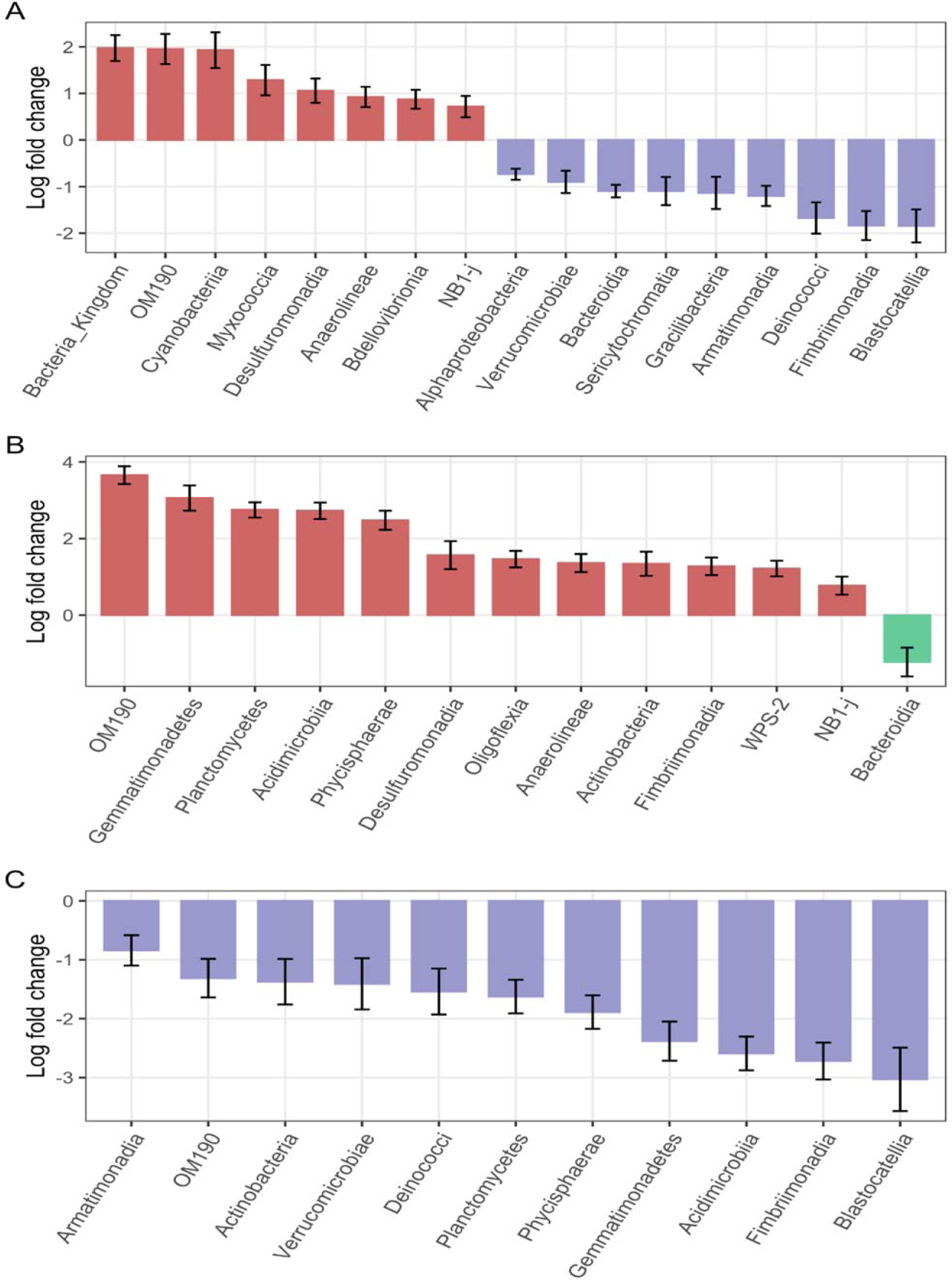
Waterfall Plots for Hatchery Surfaces. (A) Wall and baffle swabs, (B) wall and tailscreen swabs, and (C) tailscreen and baffle swabs were compared in pairwise fashion using ANCOM-BC at the class level. Log2-fold differences are plotted from classes found to be significantly different. Red bars – elevated in wall swabs, blue bars – elevated in baffle swabs, green bars – elevated in tailscreen swabs. Bacteria_Kingdom indicates reads that were not assigned past the kingdom level.

**Supplementary Figure 9:**
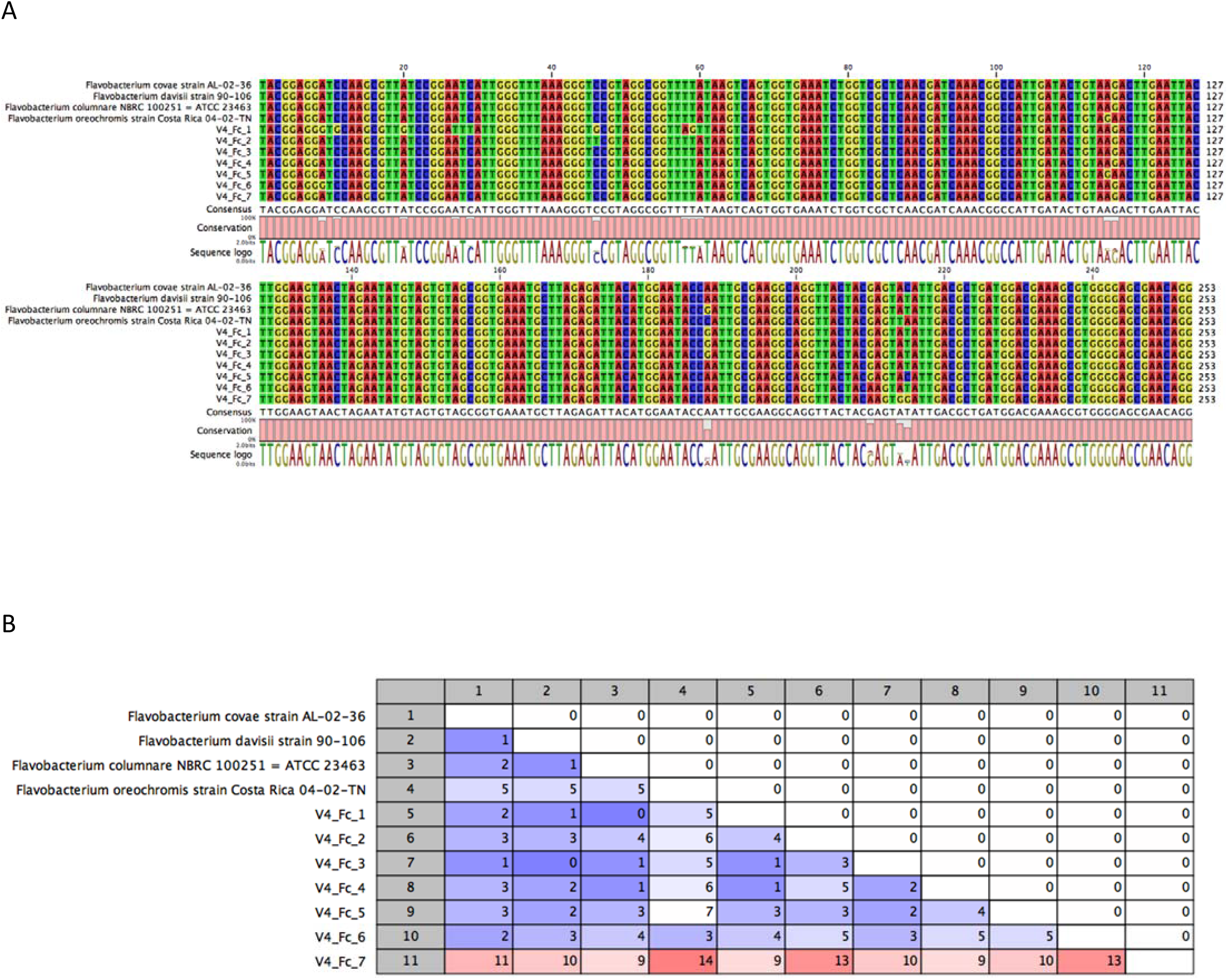
*F. columnare* 16S rRNA V4 Region Gene Sequence Comparison. 16S rRNA gene sequences from the four type strains previously comprising the single species *F. columnare* were compared with the 7 ASVs detected within the hatchery dataset. An alignment (A) and distance matrix (B) were generated for this comparison. ASVs from the hatchery dataset are in decreasing order by total read count from 1 to 7.

**Supplementary Table 1:** **Summary of Samples** ^1^Due to sample sparsity for specific sites, testing could not be performed on these interactions.

| **Sample Number** | **BioSample Accession** | **Site** | **Raceway ID** | **Year** | **Sample Type** | **Final Read Count** |
| --- | --- | --- | --- | --- | --- | --- |
| 1 | SAMN27156050 | Hatchery | 6 | 2017 | Baffle | 97335 |
| 2 | SAMN27156051 | Hatchery | 6 | 2017 | Baffle | 24975 |
| 3 | SAMN27156090 | Hatchery | 6 | 2017 | Prefilter | 36639 |
| 4 | SAMN27156117 | Hatchery | 6 | 2017 | Tailscreen | 52382 |
| 5 | SAMN27156052 | Hatchery | 8 | 2017 | Baffle | 45411 |
| 6 | SAMN27156047 | Hatchery | 14 | 2017 | Baffle | 19054 |
| 7 | SAMN27156048 | Hatchery | 19 | 2017 | Baffle | 65294 |
| 8 | SAMN27156049 | Hatchery | 20 | 2017 | Baffle | 36761 |
| 9 | SAMN27156091 | Hatchery | Global Inflow | 2017 | Prefilter | 80237 |
| 10 | SAMN27156098 | Hatchery | Global Inflow | 2017 | Sterivex | 61951 |
| 11 | SAMN27156101 | Hatchery | Global Inflow | 2017 | Sterivex | 104949 |
| 12 | SAMN27156099 | Hatchery | Global Outflow | 2017 | Sterivex | 16695 |
| 13 | SAMN27156100 | Hatchery | Global Outflow | 2017 | Sterivex | 17882 |
| 14 | SAMN27156183 | Outdoor Raceways | 11 | 2017 | Intestine | 52719 |
| 15 | SAMN27156184 | Outdoor Raceways | 11 | 2017 | Intestine | 10967 |
| 16 | SAMN27156186 | Outdoor Raceways | 11 | 2017 | Prefilter | 37641 |
| 17 | SAMN27156191 | Outdoor Raceways | 11 | 2017 | Wall | 35347 |
| 18 | SAMN27156192 | Outdoor Raceways | 11 | 2017 | Wall | 42876 |
| 19 | SAMN27156182 | Outdoor Raceways | 15 | 2017 | Gill | 14805 |
| 20 | SAMN27156181 | Outdoor Raceways | 15 | 2017 | Gill | 17319 |
| 21 | SAMN27156187 | Outdoor Raceways | 15 | 2017 | Prefilter | 39936 |
| 22 | SAMN27156193 | Outdoor Raceways | 15 | 2017 | Wall | 18868 |
| 23 | SAMN27156194 | Outdoor Raceways | 15 | 2017 | Wall | 42581 |
| 24 | SAMN27156185 | Outdoor Raceways | Global Inflow | 2017 | Prefilter | 41543 |
| 25 | SAMN27156188 | Outdoor Raceways | Global Inflow | 2017 | Sterivex | 51162 |
| 26 | SAMN27156189 | Outdoor Raceways | Global Outflow | 2017 | Sterivex | 17298 |
| 27 | SAMN27156190 | Outdoor Raceways | Global Outflow | 2017 | Sterivex | 19492 |
| 28 | SAMN27156198 | BCS | N/A | 2017 | Wall | 49281 |
| 29 | SAMN27156199 | BCS | N/A | 2017 | Wall | 46696 |
| 30 | SAMN27156056 | Hatchery | 9 | 2018 | Baffle | 100379 |
| 31 | SAMN27156057 | Hatchery | 9 | 2018 | Baffle | 53660 |
| 32 | SAMN27156058 | Hatchery | 9 | 2018 | Baffle | 121706 |
| 33 | SAMN27156059 | Hatchery | 9 | 2018 | Baffle | 35652 |
| 34 | SAMN27156092 | Hatchery | 9 | 2018 | Prefilter | 10896 |
| 35 | SAMN27156093 | Hatchery | 9 | 2018 | Prefilter | 64545 |
| 36 | SAMN27156103 | Hatchery | 9 | 2018 | Sterivex | 114452 |
| 37 | SAMN27156104 | Hatchery | 9 | 2018 | Sterivex | 55537 |
| 38 | SAMN27156119 | Hatchery | 9 | 2018 | Tailscreen | 16367 |
| 39 | SAMN27156128 | Hatchery | 9 | 2018 | Wall | 46967 |
| 40 | SAMN27156129 | Hatchery | 9 | 2018 | Wall | 47685 |
| 41 | SAMN27156130 | Hatchery | 9 | 2018 | Wall | 29580 |
| 42 | SAMN27156053 | Hatchery | 11 | 2018 | Baffle | 89277 |
| 43 | SAMN27156054 | Hatchery | 11 | 2018 | Baffle | 111266 |
| 44 | SAMN27156055 | Hatchery | 11 | 2018 | Baffle | 38021 |
| 45 | SAMN27156118 | Hatchery | 11 | 2018 | Tailscreen | 12311 |
| 46 | SAMN27156124 | Hatchery | 11 | 2018 | Wall | 123608 |
| 47 | SAMN27156125 | Hatchery | 11 | 2018 | Wall | 22624 |
| 48 | SAMN27156126 | Hatchery | 11 | 2018 | Wall | 17685 |
| 49 | SAMN27156127 | Hatchery | 16 | 2018 | Wall | 88742 |
| 50 | SAMN27156102 | Hatchery | Global Inflow | 2018 | Sterivex | 55104 |
| 51 | SAMN27156195 | BCS | N/A | 2018 | Intestine | 22981 |
| 52 | SAMN27156196 | BCS | N/A | 2018 | Intestine | 13416 |
| 53 | SAMN27156197 | BCS | N/A | 2018 | Sterivex | 86709 |
| 54 | SAMN27156200 | BCS | N/A | 2018 | Wall | 56506 |
| 55 | SAMN27156070 | Hatchery | 1 | 2019 | Baffle | 49124 |
| 56 | SAMN27156071 | Hatchery | 1 | 2019 | Baffle | 20848 |
| 57 | SAMN27156072 | Hatchery | 1 | 2019 | Baffle | 32048 |
| 58 | SAMN27156073 | Hatchery | 1 | 2019 | Baffle | 47895 |
| 59 | SAMN27156074 | Hatchery | 1 | 2019 | Baffle | 45974 |
| 60 | SAMN27156075 | Hatchery | 1 | 2019 | Baffle | 21094 |
| 61 | SAMN27156076 | Hatchery | 1 | 2019 | Baffle | 42022 |
| 62 | SAMN27156094 | Hatchery | 1 | 2019 | Prefilter | 24436 |
| 63 | SAMN27156105 | Hatchery | 1 | 2019 | Sterivex | 30241 |
| 64 | SAMN27156106 | Hatchery | 1 | 2019 | Sterivex | 27031 |
| 65 | SAMN27156160 | Hatchery | 1 | 2019 | Wall | 66874 |
| 66 | SAMN27156161 | Hatchery | 1 | 2019 | Wall | 27823 |
| 67 | SAMN27156162 | Hatchery | 1 | 2019 | Wall | 47713 |
| 68 | SAMN27156163 | Hatchery | 1 | 2019 | Wall | 31361 |
| 69 | SAMN27156164 | Hatchery | 1 | 2019 | Wall | 40100 |
| 70 | SAMN27156165 | Hatchery | 1 | 2019 | Wall | 24135 |
| 71 | SAMN27156166 | Hatchery | 1 | 2019 | Wall | 69859 |
| 72 | SAMN27156083 | Hatchery | 2 | 2019 | Baffle | 83995 |
| 73 | SAMN27156084 | Hatchery | 2 | 2019 | Baffle | 34789 |
| 74 | SAMN27156085 | Hatchery | 2 | 2019 | Baffle | 72400 |
| 75 | SAMN27156086 | Hatchery | 2 | 2019 | Baffle | 72046 |
| 76 | SAMN27156087 | Hatchery | 2 | 2019 | Baffle | 23188 |
| 77 | SAMN27156088 | Hatchery | 2 | 2019 | Baffle | 17014 |
| 78 | SAMN27156089 | Hatchery | 2 | 2019 | Baffle | 68713 |
| 79 | SAMN27156095 | Hatchery | 2 | 2019 | Prefilter | 24046 |
| 80 | SAMN27156096 | Hatchery | 2 | 2019 | Prefilter | 19179 |
| 81 | SAMN27156113 | Hatchery | 2 | 2019 | Sterivex | 15861 |
| 82 | SAMN27156114 | Hatchery | 2 | 2019 | Sterivex | 22509 |
| 83 | SAMN27156175 | Hatchery | 2 | 2019 | Wall | 46866 |
| 84 | SAMN27156176 | Hatchery | 2 | 2019 | Wall | 80912 |
| 85 | SAMN27156177 | Hatchery | 2 | 2019 | Wall | 38752 |
| 86 | SAMN27156178 | Hatchery | 2 | 2019 | Wall | 42984 |
| 87 | SAMN27156179 | Hatchery | 2 | 2019 | Wall | 24812 |
| 88 | SAMN27156180 | Hatchery | 2 | 2019 | Wall | 34524 |
| 89 | SAMN27156077 | Hatchery | 4 | 2019 | Baffle | 52165 |
| 90 | SAMN27156078 | Hatchery | 4 | 2019 | Baffle | 38246 |
| 91 | SAMN27156079 | Hatchery | 4 | 2019 | Baffle | 65713 |
| 92 | SAMN27156080 | Hatchery | 4 | 2019 | Baffle | 41429 |
| 93 | SAMN27156081 | Hatchery | 4 | 2019 | Baffle | 79989 |
| 94 | SAMN27156082 | Hatchery | 4 | 2019 | Baffle | 41698 |
| 95 | SAMN27156107 | Hatchery | 4 | 2019 | Sterivex | 48223 |
| 96 | SAMN27156108 | Hatchery | 4 | 2019 | Sterivex | 11237 |
| 97 | SAMN27156123 | Hatchery | 4 | 2019 | Tailscreen | 33234 |
| 98 | SAMN27156167 | Hatchery | 4 | 2019 | Wall | 31368 |
| 99 | SAMN27156168 | Hatchery | 4 | 2019 | Wall | 28743 |
| 100 | SAMN27156169 | Hatchery | 4 | 2019 | Wall | 44824 |
| 101 | SAMN27156170 | Hatchery | 4 | 2019 | Wall | 40921 |
| 102 | SAMN27156171 | Hatchery | 4 | 2019 | Wall | 58156 |
| 103 | SAMN27156060 | Hatchery | 8 | 2019 | Baffle | 27750 |
| 104 | SAMN27156061 | Hatchery | 8 | 2019 | Baffle | 10978 |
| 105 | SAMN27156062 | Hatchery | 8 | 2019 | Baffle | 10720 |
| 106 | SAMN27156120 | Hatchery | 8 | 2019 | Tailscreen | 52232 |
| 107 | SAMN27156121 | Hatchery | 8 | 2019 | Tailscreen | 58755 |
| 108 | SAMN27156122 | Hatchery | 8 | 2019 | Tailscreen | 26943 |
| 109 | SAMN27156131 | Hatchery | 8 | 2019 | Wall | 44277 |
| 110 | SAMN27156132 | Hatchery | 8 | 2019 | Wall | 31499 |
| 111 | SAMN27156133 | Hatchery | 8 | 2019 | Wall | 42472 |
| 112 | SAMN27156097 | Hatchery | 9 | 2019 | Prefilter | 39325 |
| 113 | SAMN27156115 | Hatchery | 9 | 2019 | Sterivex | 19687 |
| 114 | SAMN27156116 | Hatchery | 9 | 2019 | Sterivex | 32062 |
| 115 | SAMN27156172 | Hatchery | 9 | 2019 | Wall | 54288 |
| 116 | SAMN27156173 | Hatchery | 9 | 2019 | Wall | 30401 |
| 117 | SAMN27156174 | Hatchery | 9 | 2019 | Wall | 15257 |
| 118 | SAMN27156109 | Hatchery | 10 | 2019 | Sterivex | 28559 |
| 119 | SAMN27156110 | Hatchery | 10 | 2019 | Sterivex | 25746 |
| 120 | SAMN27156134 | Hatchery | 10 | 2019 | Wall | 30488 |
| 121 | SAMN27156135 | Hatchery | 10 | 2019 | Wall | 55582 |
| 122 | SAMN27156136 | Hatchery | 10 | 2019 | Wall | 65335 |
| 123 | SAMN27156137 | Hatchery | 10 | 2019 | Wall | 83071 |
| 124 | SAMN27156138 | Hatchery | 11 | 2019 | Wall | 18941 |
| 125 | SAMN27156139 | Hatchery | 11 | 2019 | Wall | 41404 |
| 126 | SAMN27156140 | Hatchery | 11 | 2019 | Wall | 40264 |
| 127 | SAMN27156141 | Hatchery | 11 | 2019 | Wall | 37745 |
| 128 | SAMN27156063 | Hatchery | 12 | 2019 | Baffle | 38024 |
| 129 | SAMN27156064 | Hatchery | 12 | 2019 | Baffle | 41947 |
| 130 | SAMN27156065 | Hatchery | 12 | 2019 | Baffle | 53911 |
| 131 | SAMN27156066 | Hatchery | 12 | 2019 | Baffle | 47991 |
| 132 | SAMN27156067 | Hatchery | 12 | 2019 | Baffle | 55628 |
| 133 | SAMN27156068 | Hatchery | 12 | 2019 | Baffle | 118206 |
| 134 | SAMN27156069 | Hatchery | 12 | 2019 | Baffle | 80312 |
| 135 | SAMN27156142 | Hatchery | 12 | 2019 | Wall | 50708 |
| 136 | SAMN27156143 | Hatchery | 12 | 2019 | Wall | 50835 |
| 137 | SAMN27156144 | Hatchery | 12 | 2019 | Wall | 172746 |
| 138 | SAMN27156145 | Hatchery | 12 | 2019 | Wall | 164648 |
| 139 | SAMN27156111 | Hatchery | 16 | 2019 | Sterivex | 26260 |
| 140 | SAMN27156112 | Hatchery | 16 | 2019 | Sterivex | 26428 |
| 141 | SAMN27156146 | Hatchery | 16 | 2019 | Wall | 30427 |
| 142 | SAMN27156147 | Hatchery | 16 | 2019 | Wall | 24192 |
| 143 | SAMN27156148 | Hatchery | 16 | 2019 | Wall | 34118 |
| 144 | SAMN27156149 | Hatchery | 16 | 2019 | Wall | 21829 |
| 145 | SAMN27156150 | Hatchery | 17 | 2019 | Wall | 33660 |
| 146 | SAMN27156151 | Hatchery | 17 | 2019 | Wall | 45161 |
| 147 | SAMN27156152 | Hatchery | 17 | 2019 | Wall | 16197 |
| 148 | SAMN27156153 | Hatchery | 17 | 2019 | Wall | 23562 |
| 149 | SAMN27156154 | Hatchery | 18 | 2019 | Wall | 22460 |
| 150 | SAMN27156155 | Hatchery | 18 | 2019 | Wall | 50796 |
| 151 | SAMN27156156 | Hatchery | 18 | 2019 | Wall | 16011 |
| 152 | SAMN27156157 | Hatchery | 18 | 2019 | Wall | 52423 |
| 153 | SAMN27156158 | Hatchery | 18 | 2019 | Wall | 31999 |
| 154 | SAMN27156159 | Hatchery | 18 | 2019 | Wall | 30377 |
| 155 | SAMN27156201 | BCS | N/A | 2019 | Wall | 46605 |
| 156 | SAMN27156202 | BCS | N/A | 2019 | Wall | 12823 |
| 157 | SAMN27156203 | BCS | N/A | 2019 | Wall | 76585 |
| 158 | SAMN27156204 | BCS | N/A | 2019 | Wall | 42970 |
| 159 | SAMN27156205 | BCS | N/A | 2019 | Wall | 87642 |
| 160 | SAMN27156206 | BCS | N/A | 2019 | Wall | 95144 |
| 161 | SAMN27156207 | BCS | N/A | 2019 | Wall | 42303 |
| 162 | SAMN27156208 | BCS | N/A | 2019 | Wall | 78732 |
| 163 | SAMN27156209 | BCS | N/A | 2019 | Wall | 42722 |

**Supplementary Table 2:** PERMANOVA Testing of Bray-Curtis Distances for Surface and Water Samples at the Three Sites.

|  | Surfaces |  | Water |  |
| --- | --- | --- | --- | --- |
|  | p | R<sup>2</sup> | p | R<sup>2</sup> |
| Sample Type | 0.0001 | 0.0909 | 0.0001 | 0.3117 |
| Sampling Year | 0.0001 | 0.0328 | 0.0186 | 0.0476 |
| Sampling Site | 0.0001 | 0.0882 | 0.1408 | 0.0517 |
| Sample Type: Year | 0.0001 | 0.0351 | 0.1421 | 0.0268 |
| Sampling Site: Year<sup>1</sup> | 0.0001 | 0.0217 |  |  |
| Sample Type: Site<sup>1</sup> |  |  | 0.3472 | 0.0193 |

<sup>1</sup>Due to sample sparsity for specific sites, testing could not be performed on these interactions

**Supplementary Table 3:**
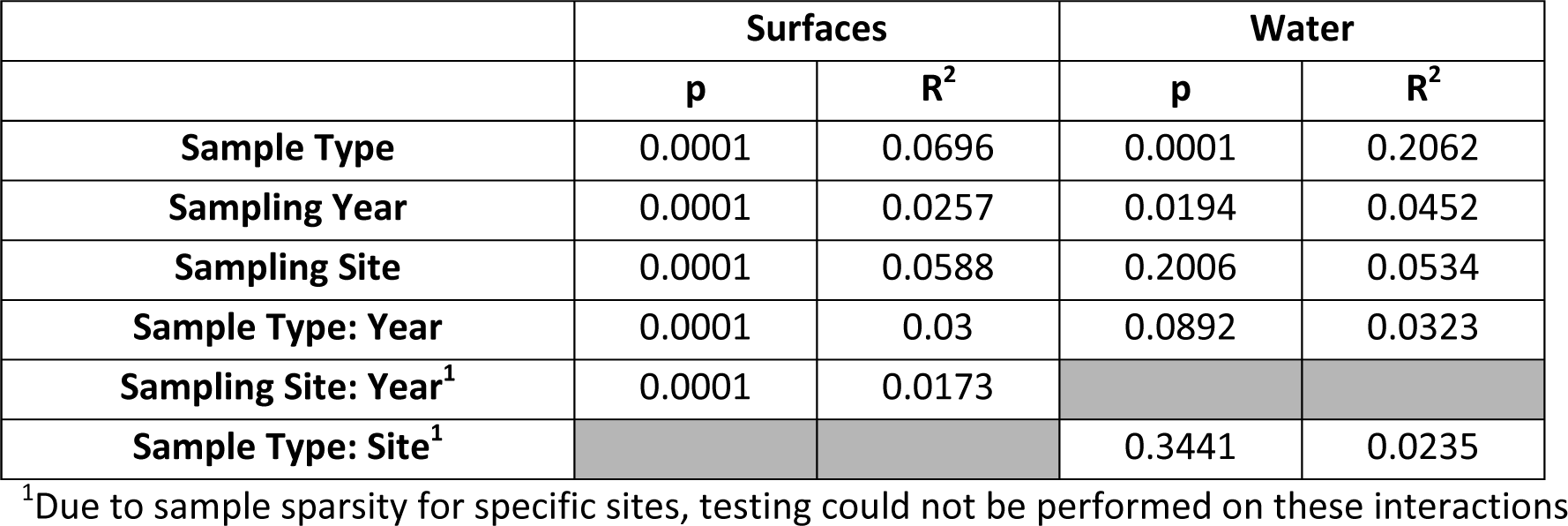
PERMANOVA Testing of Jaccard Distances for Surface and Water Samples at the Three Sites.

**Supplementary Table 4:** PERMANOVA Testing of Unweighted Unifrac Distances for Surface and Water Samples at Three Sites.

|  | Surfaces |  | Water |  |
| --- | --- | --- | --- | --- |
|  | p | R<sup>2</sup> | p | R<sup>2</sup> |
| Sample Type | 0.0001 | 0.0894 | 0.0001 | 0.2598 |
| Sampling Year | 0.0001 | 0.0232 | 0.2348 | 0.0254 |
| Sampling Site | 0.0001 | 0.0723 | 0.3346 | 0.0463 |
| Sample Type: Year | 0.0002 | 0.0282 | 0.3479 | 0.0221 |
| Sampling Site: Year<sup>1</sup> | 0.0006 | 0.0163 |  |  |
| Sample Type: Site<sup>1</sup> |  |  | 0.3877 | 0.0214 |

<sup>1</sup>Due to sample sparsity for specific sites, testing could not be performed on these interactions

**Supplementary Table 5:**
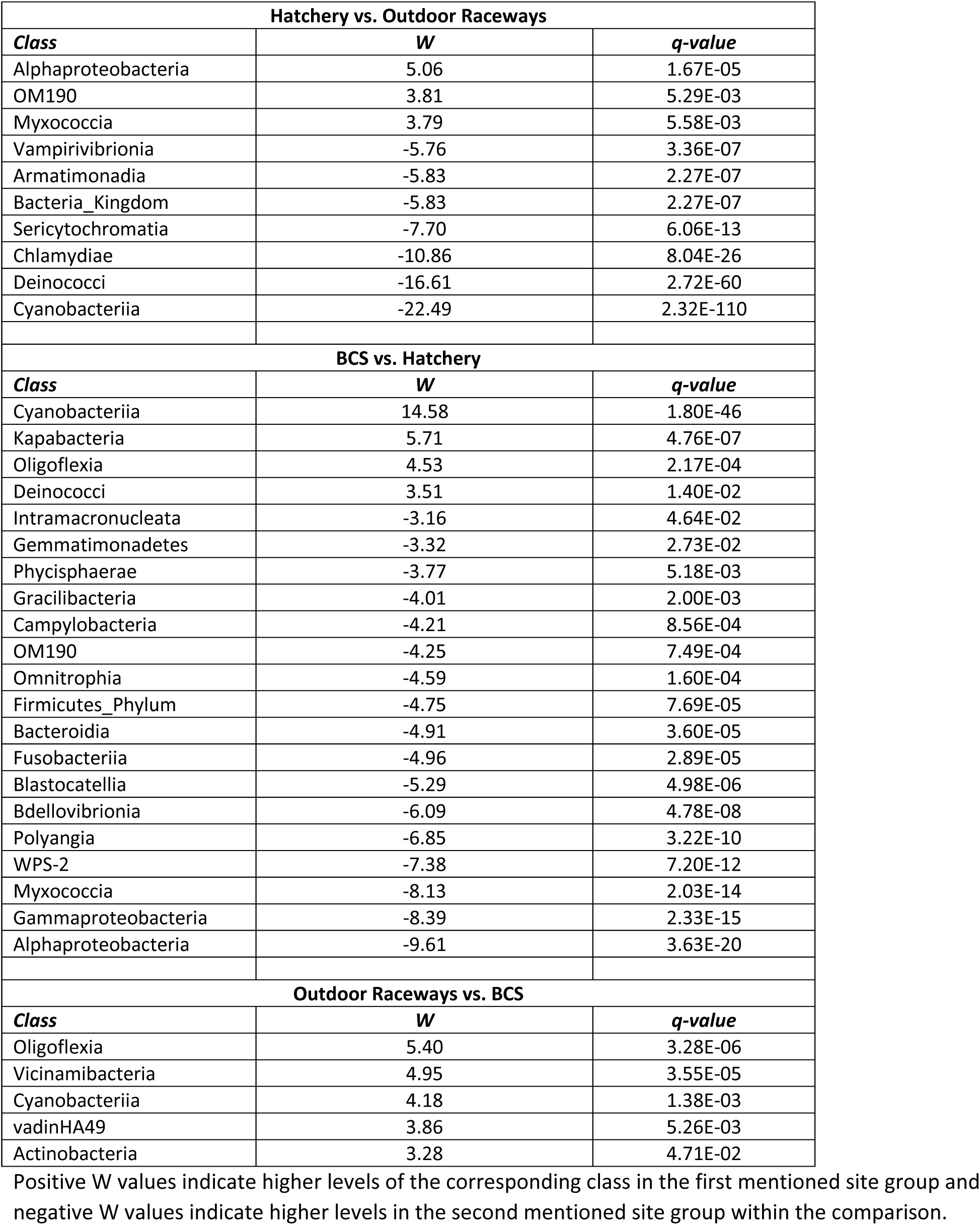
Significant Class-Level ANCOM Results Between Sampling Sites.

**Supplementary Table 6:**
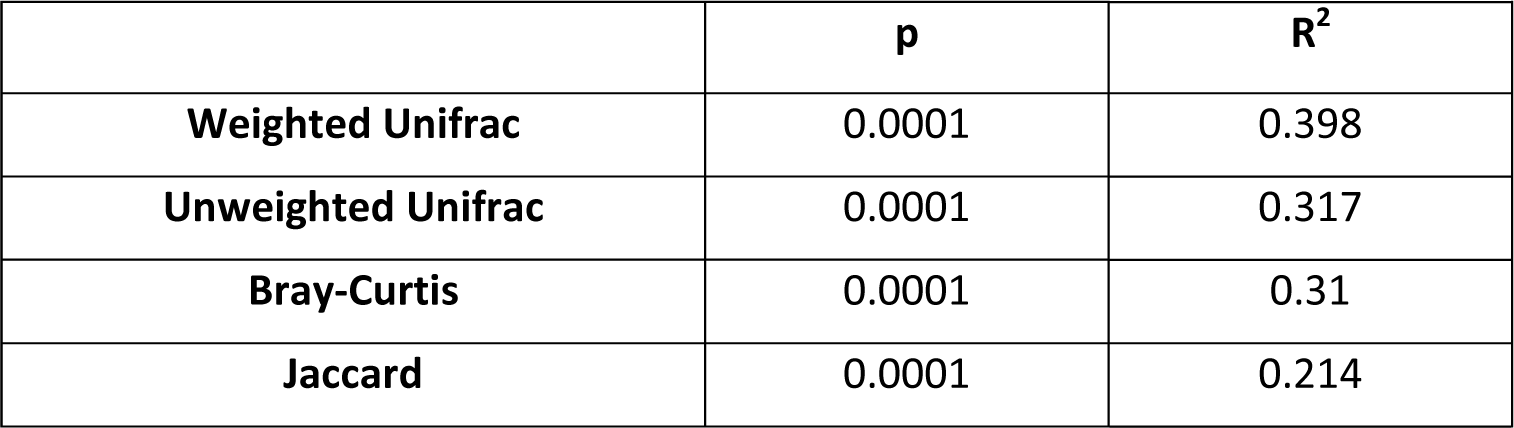
PERMANOVA Testing Results of Sample Type Variable within Hatchery Samples for All Methods.

**Supplementary Table 7:**
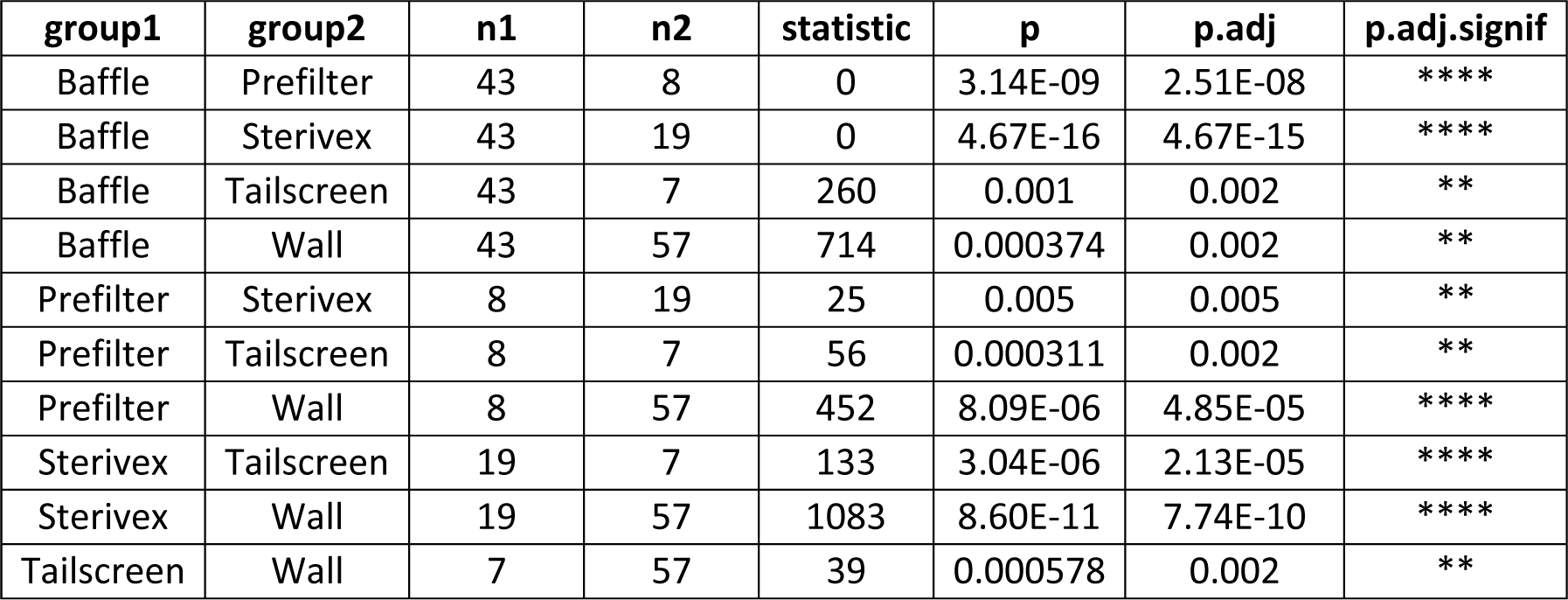
Pairwise Wilcoxon Signed-Rank Test Statistics for Shannon Index Diversity by Sample Type.

| **group1** | **group2** | **n1** | **n2** | **statistic** | **p** | **p.adj** | **p.adj.signif** |
| --- | --- | --- | --- | --- | --- | --- | --- |
| Baffle | Prefilter | 43 | 8 | 0 | 3.14E-09 | 2.51E-08 | **** |
| Baffle | Sterivex | 43 | 19 | 0 | 4.67E-16 | 4.67E-15 | **** |
| Baffle | Tailscreen | 43 | 7 | 260 | 0.001 | 0.002 | ** |
| Baffle | Wall | 43 | 57 | 714 | 0.000374 | 0.002 | ** |
| Prefilter | Sterivex | 8 | 19 | 25 | 0.005 | 0.005 | ** |
| Prefilter | Tailscreen | 8 | 7 | 56 | 0.000311 | 0.002 | ** |
| Prefilter | Wall | 8 | 57 | 452 | 8.09E-06 | 4.85E-05 | **** |
| Sterivex | Tailscreen | 19 | 7 | 133 | 3.04E-06 | 2.13E-05 | **** |
| Sterivex | Wall | 19 | 57 | 1083 | 8.60E-11 | 7.74E-10 | **** |
| Tailscreen | Wall | 7 | 57 | 39 | 0.000578 | 0.002 | ** |

**Supplementary Table 8:** Pairwise Wilcoxon Signed-Rank Test Statistics for Faith’s Phylogenetic Diversity by Sample Type.

| **group1** | **group2** | **n1** | **n2** | **statistic** | **p** | **p.adj** | **p.adj.signif** |
| --- | --- | --- | --- | --- | --- | --- | --- |
| Baffle | Prefilter | 43 | 8 | 1 | 6.28E-09 | 5.02E-08 | **** |
| Baffle | Sterivex | 43 | 19 | 0 | 4.67E-16 | 4.67E-15 | **** |
| Baffle | Tailscreen | 43 | 7 | 219 | 0.056 | 0.059 | ns |
| Baffle | Wall | 43 | 57 | 534 | 1.50E-06 | 1.05E-05 | **** |
| Prefilter | Sterivex | 8 | 19 | 35 | 0.029 | 0.059 | ns |
| Prefilter | Tailscreen | 8 | 7 | 56 | 0.000311 | 0.000933 | *** |
| Prefilter | Wall | 8 | 57 | 454 | 6.71E-06 | 3.36E-05 | **** |
| Sterivex | Tailscreen | 19 | 7 | 133 | 3.04E-06 | 1.82E-05 | **** |
| Sterivex | Wall | 19 | 57 | 1083 | 8.60E-11 | 7.74E-10 | **** |
| Tailscreen | Wall | 7 | 57 | 27 | 0.000216 | 0.000864 | *** |

**Supplementary Table 9:**
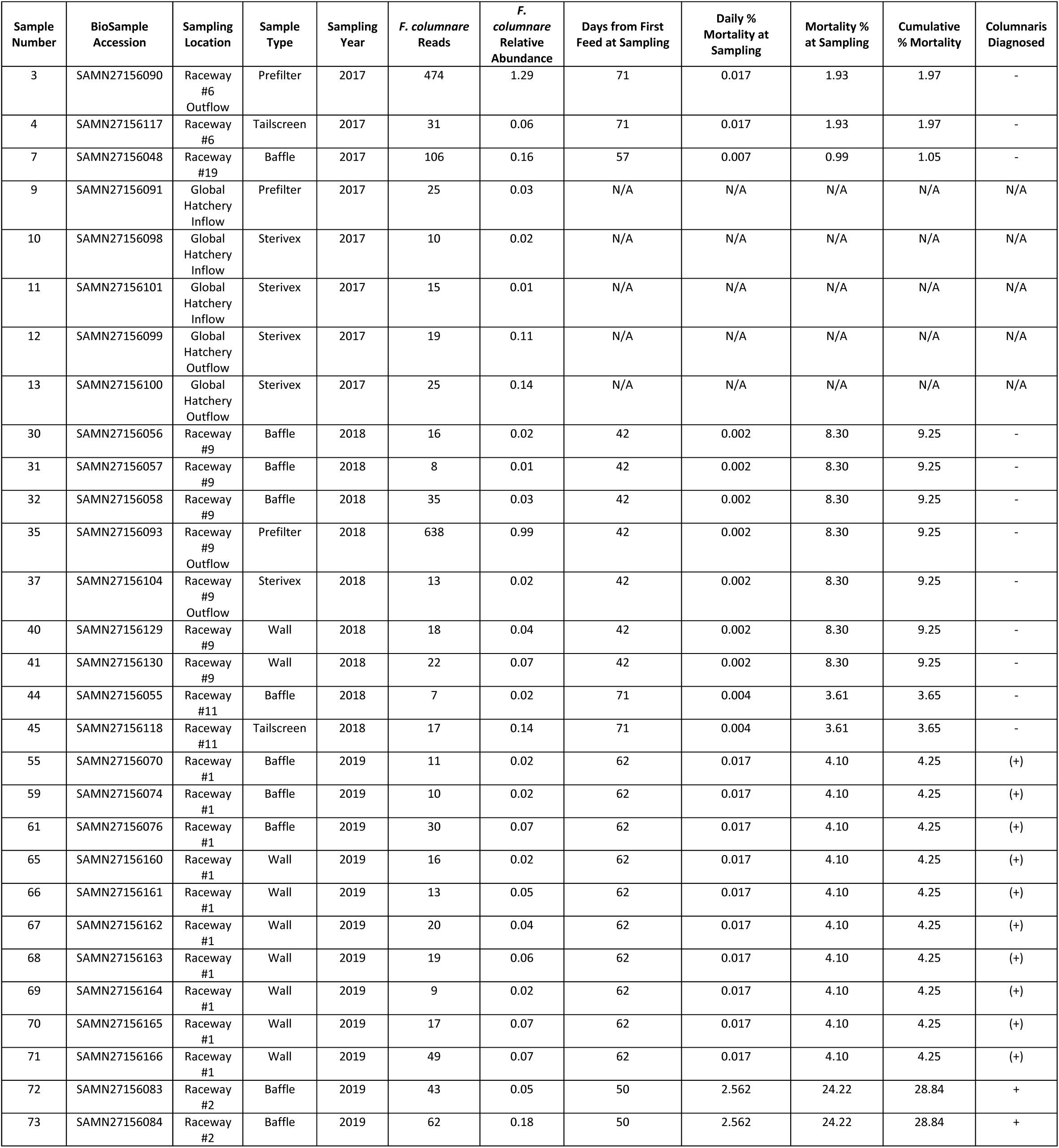
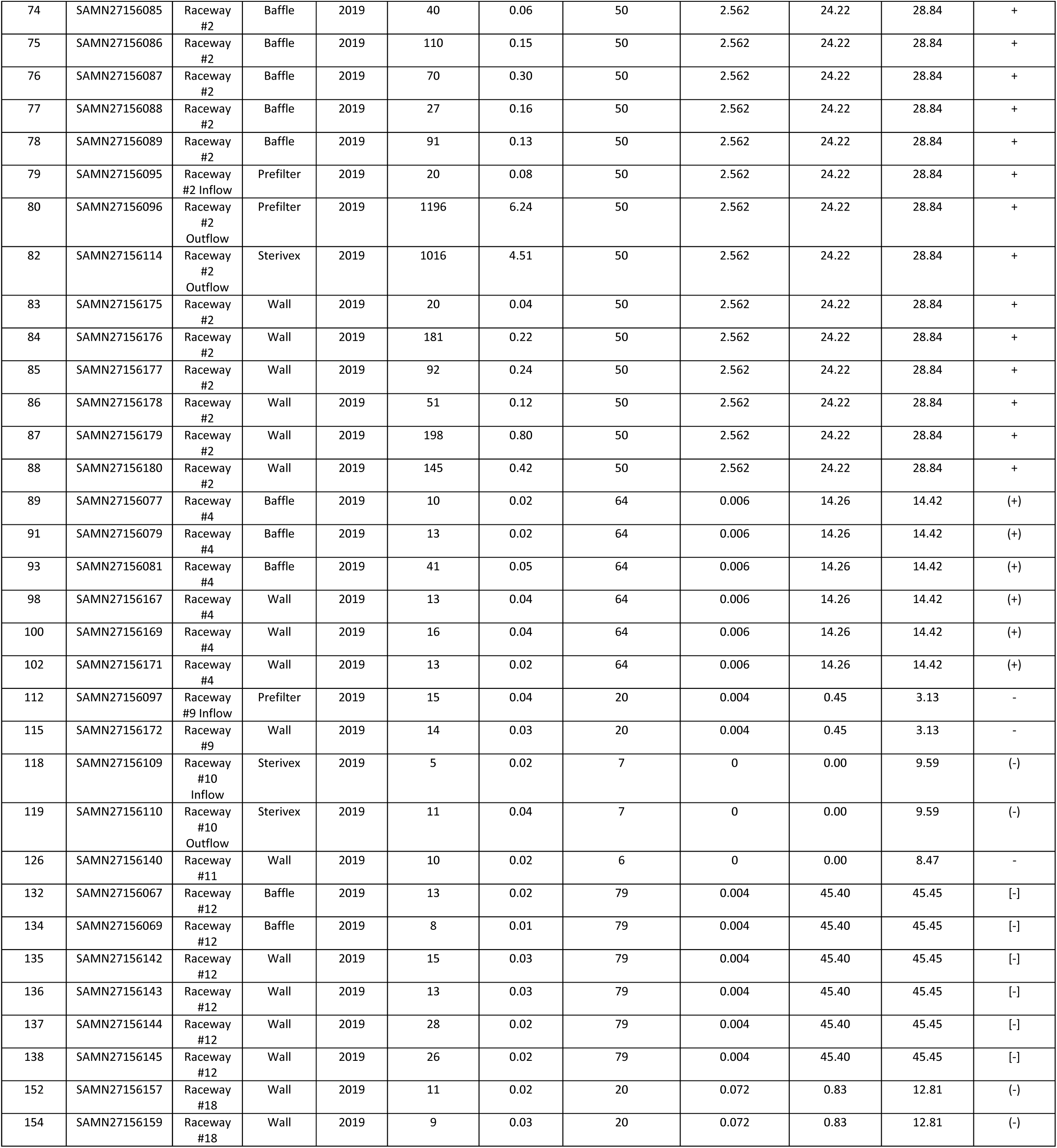
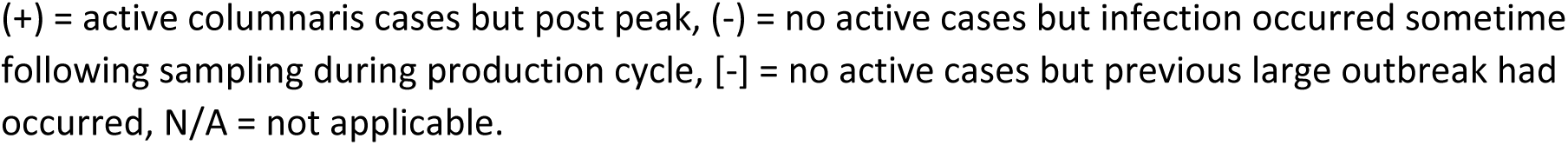
*F. columnare*-positive Samples.

| Sample Number | BioSample Accession | Sampling Location | Sample Type | Sampling Year | *F. columnare* Reads | *F. columnare* Relative Abundance | Days from First Feed at Sampling | Daily % Mortality at Sampling | Mortality % at Sampling | Cumulative % Mortality | Columnaris Diagnosed |
| --- | --- | --- | --- | --- | --- | --- | --- | --- | --- | --- | --- |
| 3 | SAMN27156090 | Raceway #6 Outflow | Prefilter | 2017 | 474 | 1.29 | 71 | 0.017 | 1.93 | 1.97 | - |
| 4 | SAMN27156117 | Raceway #6 | Tailscreen | 2017 | 31 | 0.06 | 71 | 0.017 | 1.93 | 1.97 | - |
| 7 | SAMN27156048 | Raceway #19 | Baffle | 2017 | 106 | 0.16 | 57 | 0.007 | 0.99 | 1.05 | - |
| 9 | SAMN27156091 | Global Hatchery Inflow | Prefilter | 2017 | 25 | 0.03 | N/A | N/A | N/A | N/A | N/A |
| 10 | SAMN27156098 | Global Hatchery Inflow | Sterivex | 2017 | 10 | 0.02 | N/A | N/A | N/A | N/A | N/A |
| 11 | SAMN27156101 | Global Hatchery Inflow | Sterivex | 2017 | 15 | 0.01 | N/A | N/A | N/A | N/A | N/A |
| 12 | SAMN27156099 | Global Hatchery Outflow | Sterivex | 2017 | 19 | 0.11 | N/A | N/A | N/A | N/A | N/A |
| 13 | SAMN27156100 | Global Hatchery Outflow | Sterivex | 2017 | 25 | 0.14 | N/A | N/A | N/A | N/A | N/A |
| 30 | SAMN27156056 | Raceway #9 | Baffle | 2018 | 16 | 0.02 | 42 | 0.002 | 8.30 | 9.25 | - |
| 31 | SAMN27156057 | Raceway #9 | Baffle | 2018 | 8 | 0.01 | 42 | 0.002 | 8.30 | 9.25 | - |
| 32 | SAMN27156058 | Raceway #9 | Baffle | 2018 | 35 | 0.03 | 42 | 0.002 | 8.30 | 9.25 | - |
| 35 | SAMN27156093 | Raceway #9 Outflow | Prefilter | 2018 | 638 | 0.99 | 42 | 0.002 | 8.30 | 9.25 | - |
| 37 | SAMN27156104 | Raceway #9 Outflow | Sterivex | 2018 | 13 | 0.02 | 42 | 0.002 | 8.30 | 9.25 | - |
| 40 | SAMN27156129 | Raceway #9 | Wall | 2018 | 18 | 0.04 | 42 | 0.002 | 8.30 | 9.25 | - |
| 41 | SAMN27156130 | Raceway #9 | Wall | 2018 | 22 | 0.07 | 42 | 0.002 | 8.30 | 9.25 | - |
| 44 | SAMN27156055 | Raceway #11 | Baffle | 2018 | 7 | 0.02 | 71 | 0.004 | 3.61 | 3.65 | - |
| 45 | SAMN27156118 | Raceway #11 | Tailscreen | 2018 | 17 | 0.14 | 71 | 0.004 | 3.61 | 3.65 | - |
| 55 | SAMN27156070 | Raceway #1 | Baffle | 2019 | 11 | 0.02 | 62 | 0.017 | 4.10 | 4.25 | (+) |
| 59 | SAMN27156074 | Raceway #1 | Baffle | 2019 | 10 | 0.02 | 62 | 0.017 | 4.10 | 4.25 | (+) |
| 61 | SAMN27156076 | Raceway #1 | Baffle | 2019 | 30 | 0.07 | 62 | 0.017 | 4.10 | 4.25 | (+) |
| 65 | SAMN27156160 | Raceway #1 | Wall | 2019 | 16 | 0.02 | 62 | 0.017 | 4.10 | 4.25 | (+) |
| 66 | SAMN27156161 | Raceway #1 | Wall | 2019 | 13 | 0.05 | 62 | 0.017 | 4.10 | 4.25 | (+) |
| 67 | SAMN27156162 | Raceway #1 | Wall | 2019 | 20 | 0.04 | 62 | 0.017 | 4.10 | 4.25 | (+) |
| 68 | SAMN27156163 | Raceway #1 | Wall | 2019 | 19 | 0.06 | 62 | 0.017 | 4.10 | 4.25 | (+) |
| 69 | SAMN27156164 | Raceway #1 | Wall | 2019 | 9 | 0.02 | 62 | 0.017 | 4.10 | 4.25 | (+) |
| 70 | SAMN27156165 | Raceway #1 | Wall | 2019 | 17 | 0.07 | 62 | 0.017 | 4.10 | 4.25 | (+) |
| 71 | SAMN27156166 | Raceway #1 | Wall | 2019 | 49 | 0.07 | 62 | 0.017 | 4.10 | 4.25 | (+) |
| 72 | SAMN27156083 | Raceway #2 | Baffle | 2019 | 43 | 0.05 | 50 | 2.562 | 24.22 | 28.84 | + |
| 73 | SAMN27156084 | Raceway #2 | Baffle | 2019 | 62 | 0.18 | 50 | 2.562 | 24.22 | 28.84 | + |
| 74 | SAMN27156085 | Raceway #2 | Baffle | 2019 | 40 | 0.06 | 50 | 2.562 | 24.22 | 28.84 | + |
| 75 | SAMN27156086 | Raceway #2 | Baffle | 2019 | 110 | 0.15 | 50 | 2.562 | 24.22 | 28.84 | + |
| 76 | SAMN27156087 | Raceway #2 | Baffle | 2019 | 70 | 0.30 | 50 | 2.562 | 24.22 | 28.84 | + |
| 77 | SAMN27156088 | Raceway #2 | Baffle | 2019 | 27 | 0.16 | 50 | 2.562 | 24.22 | 28.84 | + |
| 78 | SAMN27156089 | Raceway #2 | Baffle | 2019 | 91 | 0.13 | 50 | 2.562 | 24.22 | 28.84 | + |
| 79 | SAMN27156095 | Raceway #2 Inflow | Prefilter | 2019 | 20 | 0.08 | 50 | 2.562 | 24.22 | 28.84 | + |
| 80 | SAMN27156096 | Raceway #2 Outflow | Prefilter | 2019 | 1196 | 6.24 | 50 | 2.562 | 24.22 | 28.84 | + |
| 82 | SAMN27156114 | Raceway #2 Outflow | Sterivex | 2019 | 1016 | 4.51 | 50 | 2.562 | 24.22 | 28.84 | + |
| 83 | SAMN27156175 | Raceway #2 | Wall | 2019 | 20 | 0.04 | 50 | 2.562 | 24.22 | 28.84 | + |
| 84 | SAMN27156176 | Raceway #2 | Wall | 2019 | 181 | 0.22 | 50 | 2.562 | 24.22 | 28.84 | + |
| 85 | SAMN27156177 | Raceway #2 | Wall | 2019 | 92 | 0.24 | 50 | 2.562 | 24.22 | 28.84 | + |
| 86 | SAMN27156178 | Raceway #2 | Wall | 2019 | 51 | 0.12 | 50 | 2.562 | 24.22 | 28.84 | + |
| 87 | SAMN27156179 | Raceway #2 | Wall | 2019 | 198 | 0.80 | 50 | 2.562 | 24.22 | 28.84 | + |
| 88 | SAMN27156180 | Raceway #2 | Wall | 2019 | 145 | 0.42 | 50 | 2.562 | 24.22 | 28.84 | + |
| 89 | SAMN27156077 | Raceway #4 | Baffle | 2019 | 10 | 0.02 | 64 | 0.006 | 14.26 | 14.42 | (+) |
| 91 | SAMN27156079 | Raceway #4 | Baffle | 2019 | 13 | 0.02 | 64 | 0.006 | 14.26 | 14.42 | (+) |
| 93 | SAMN27156081 | Raceway #4 | Baffle | 2019 | 41 | 0.05 | 64 | 0.006 | 14.26 | 14.42 | (+) |
| 98 | SAMN27156167 | Raceway #4 | Wall | 2019 | 13 | 0.04 | 64 | 0.006 | 14.26 | 14.42 | (+) |
| 100 | SAMN27156169 | Raceway #4 | Wall | 2019 | 16 | 0.04 | 64 | 0.006 | 14.26 | 14.42 | (+) |
| 102 | SAMN27156171 | Raceway #4 | Wall | 2019 | 13 | 0.02 | 64 | 0.006 | 14.26 | 14.42 | (+) |
| 112 | SAMN27156097 | Raceway #9 Inflow | Prefilter | 2019 | 15 | 0.04 | 20 | 0.004 | 0.45 | 3.13 | - |
| 115 | SAMN27156172 | Raceway #9 | Wall | 2019 | 14 | 0.03 | 20 | 0.004 | 0.45 | 3.13 | - |
| 118 | SAMN27156109 | Raceway #10 Inflow | Sterivex | 2019 | 5 | 0.02 | 7 | 0 | 0.00 | 9.59 | (-) |
| 119 | SAMN27156110 | Raceway #10 Outflow | Sterivex | 2019 | 11 | 0.04 | 7 | 0 | 0.00 | 9.59 | (-) |
| 126 | SAMN27156140 | Raceway #11 | Wall | 2019 | 10 | 0.02 | 6 | 0 | 0.00 | 8.47 | - |
| 132 | SAMN27156067 | Raceway #12 | Baffle | 2019 | 13 | 0.02 | 79 | 0.004 | 45.40 | 45.45 | [-] |
| 134 | SAMN27156069 | Raceway #12 | Baffle | 2019 | 8 | 0.01 | 79 | 0.004 | 45.40 | 45.45 | [-] |
| 135 | SAMN27156142 | Raceway #12 | Wall | 2019 | 15 | 0.03 | 79 | 0.004 | 45.40 | 45.45 | [-] |
| 136 | SAMN27156143 | Raceway #12 | Wall | 2019 | 13 | 0.03 | 79 | 0.004 | 45.40 | 45.45 | [-] |
| 137 | SAMN27156144 | Raceway #12 | Wall | 2019 | 28 | 0.02 | 79 | 0.004 | 45.40 | 45.45 | [-] |
| 138 | SAMN27156145 | Raceway #12 | Wall | 2019 | 26 | 0.02 | 79 | 0.004 | 45.40 | 45.45 | [-] |
| 152 | SAMN27156157 | Raceway #18 | Wall | 2019 | 11 | 0.02 | 20 | 0.072 | 0.83 | 12.81 | (-) |
| 154 | SAMN27156159 | Raceway #18 | Wall | 2019 | 9 | 0.03 | 20 | 0.072 | 0.83 | 12.81 | (-) |

(+) = active columnaris cases but post peak, (-) = no active cases but infection occurred sometime following sampling during production cycle, [-] = no active cases but previous large outbreak had occurred, N/A = not applicable.

